# Optimized dCas9 Programmable Transcriptional Activators for Plants

**DOI:** 10.1101/2022.06.10.495638

**Authors:** Matthew H Zinselmeier, J. Armando Casas-Mollano, Jonathan Cors, Adam Sychla, Stephen C Heinsch, Daniel F Voytas, Michael J Smanski

**Affiliations:** Department of Genetics, Cell Biology, and Development, University of Minnesota, Minneapolis, MN 55455; Department of Biochemistry, Molecular Biology, and Biophysics, University of Minnesota, Minneapolis, MN, 55455; Center for Precision Plant Genomics, 1500 Gortner Avenue, Cargill Building, Saint Paul, MN 55108; Biotechnology Institute, University of Minnesota, Saint Paul, MN 55108; Bioinformatics and Computational Biology Graduate Program, University of Minnesota, Saint Paul, MN 55108

**Keywords:** dCas9, Transcriptional activator, Plant engineering, SunTag

## Abstract

Understanding how gene expression impacts plant development and physiology is important for rationally engineering improved crops. Programmable Transcriptional Activators (PTAs), including CRISPR-Cas activators, have traditionally relied on a limited number of transcriptional activation domains. Usually the VP64 domain, derived from human herpes simplex virus, is fused to a DNA-binding domain to activate target gene expression. We reasoned there was considerable space for PTA improvement by replacing VP64 with a plant-derived activation domain. To address this, we designed, built, and tested a PTA library of 38 putative plant transcriptional activation domains. Domains from *HSFA6b, AvrXa10, DOF1, DREB1*, and *DREB2* genes function as strong activators in *Setaria viridis* and *Arabidopsis thaliana* in protoplast assays and in transgenic plants. Overexpression of multiple endogenous genes (*FT, PAP1, WUS*) reached levels similar to the highly expressed housekeeping gene, *PP2A*, regardless of basal expression level. Further, these domains were effective in different PTA architectures, including the dCas9-SunTag, dCas9-Moontag, and TALE-SunTag systems. Lastly, we demonstrate the ability of these improved PTAs to map enhancer regions that promote gene expression in plants. This work demonstrates the effective and flexible nature of PTAs to activate target genes in plants, providing tools that can be used to improve agronomically relevant traits of interest.

## Introduction

Controlling the expression of endogenous genes in plants is important for basic research on plant development. Genomes provide the blueprint for plant growth, survival, and reproduction. In order to survive a plant must also respond to surrounding environmental conditions such as temperature, light, or humidity. Gene expression is the process by which the genetic blueprint is put into action; activating genes at different times and places to develop, survive, and reproduce in the face of external stimuli (1, 2). Studies correlating differential gene expression with phenotype allow for predicting which genes are most important for proper development and responding to the environment (3, 4). The ability to control the spatio-temporal expression of these key genes driving phenotypic variation will be critical for trait and yield improvement of crop species in different environments across the world.

To this end, recent advances have led to the development of tools capable of activating gene expression in a user-controlled manner. Programmable Transcriptional Activators (PTAs) are fusion proteins comprised of a transcriptional activation domain (AD) fused to a DNA-binding domain that can be rationally engineered to recognize a DNA sequence of interest. The DNA-binding domain can be a zinc-finger (5), a transcriptional activator-like effector (TALE) DNA-binding domain (6), or a catalytically ‘dead’ Cas9 protein (dCas9) that is directed to a target sequence via a single guide RNA (sgRNA) (7). PTAs can drive the over- and/or ectopic-expression of endogenous genes when designed to bind to a promoter region (8, 9). The strength of overexpression is inversely correlated to a target gene’s basal expression levels; PTAs targeting lowly-expressed genes can achieve higher fold-overexpression values than targeting highly-expressed genes (10). dCas9-based PTAs can also target many promoters in parallel by co-expressing multiple sgRNAs (11). Further, PTAs have been used to explore gene function and engineer barriers to sexual reproduction in insects (12).

The first dCas9-based PTA fused the VP64 AD the dCas9 DNA binding domain. VP64 is a tetrameric repeat of the VP16 protein derived from herpes simplex virus (13, 14). This dCas9-VP64 design has been used in plants to activate target genes (15). An additional improvement to PTAs came with the dCas9 translational fusion to the VPR AD (16). VPR is a translational fusion of three different ADs - VP64 from the Herpes Simplex Virus VP16 protein, RTA from the *Homo sapien RELA* transcription factor, and p65 from the Epstein-Barr virus *BRLF1* gene. While VPR drives strong gene expression in animal species, the same robust effectiveness seen in mammalian cells remains elusive in plants (7).

Several groups have developed improved dCas9-based PTAs for use in plant systems. The most common strategy is to increase the number of ADs recruited to the PTA. dCas9-TV is comprised of 6 copies of the TAL activation domain and two copies of the VP64 activation domain translationally fused to dCas9 (17). The SunTag system uses a non-covalent interaction between single-chain variable fragment antibodies (scFv) and a tandemly-repeated epitope tail (GCN4 motif) to recruit ADs to the dCas9 (18). Using 10 copies of GCN4 fused to dCas9, the SunTag system can activate target genes in *A. thaliana* to produce visible phenotypes (19). Subsequent iterations of the SunTag system have proven highly effective in activating target genes across monocot and dicot species (20). While these PTAs improved performance in plants, many still rely on the VP64 AD. Given the evolutionary distance between mammals and plants, we sought to improve PTA activity by screening ADs that evolved in plant systems. Here we report the performance of several plant-derived ADs for programmable gene expression in monocot and dicot plants.

## Results

### Building a library of putative plant-derived transcriptional activation domains

We first determined a suitable experimental system for identifying strong ADs. Protoplast isolation and transformation pipelines yield consistent cell numbers and transformation efficiencies, allowing for direct comparison of groups within a given transformation when activating a reporter gene (21, 22).

We compared the ability of the direct-fusion dCas9-VP64 PTA architecture with the TAD-scaffolding PTA architecture SunTag (Figure 1a-b) to activate an endogenous target gene in *A. thaliana*. A tRNA-based multi-guide expression array (23) expressing four sgRNAs targeting a single core promoter for *WUSCHEL (WUS)* was transformed into *A. thaliana* protoplasts along with a dCas9 PTA, followed by RNA extraction and RT-qPCR. The dCas9-PTAs were expressed using the *Ubi10* promoter from *A. thaliana*.

**Fig. 1.**
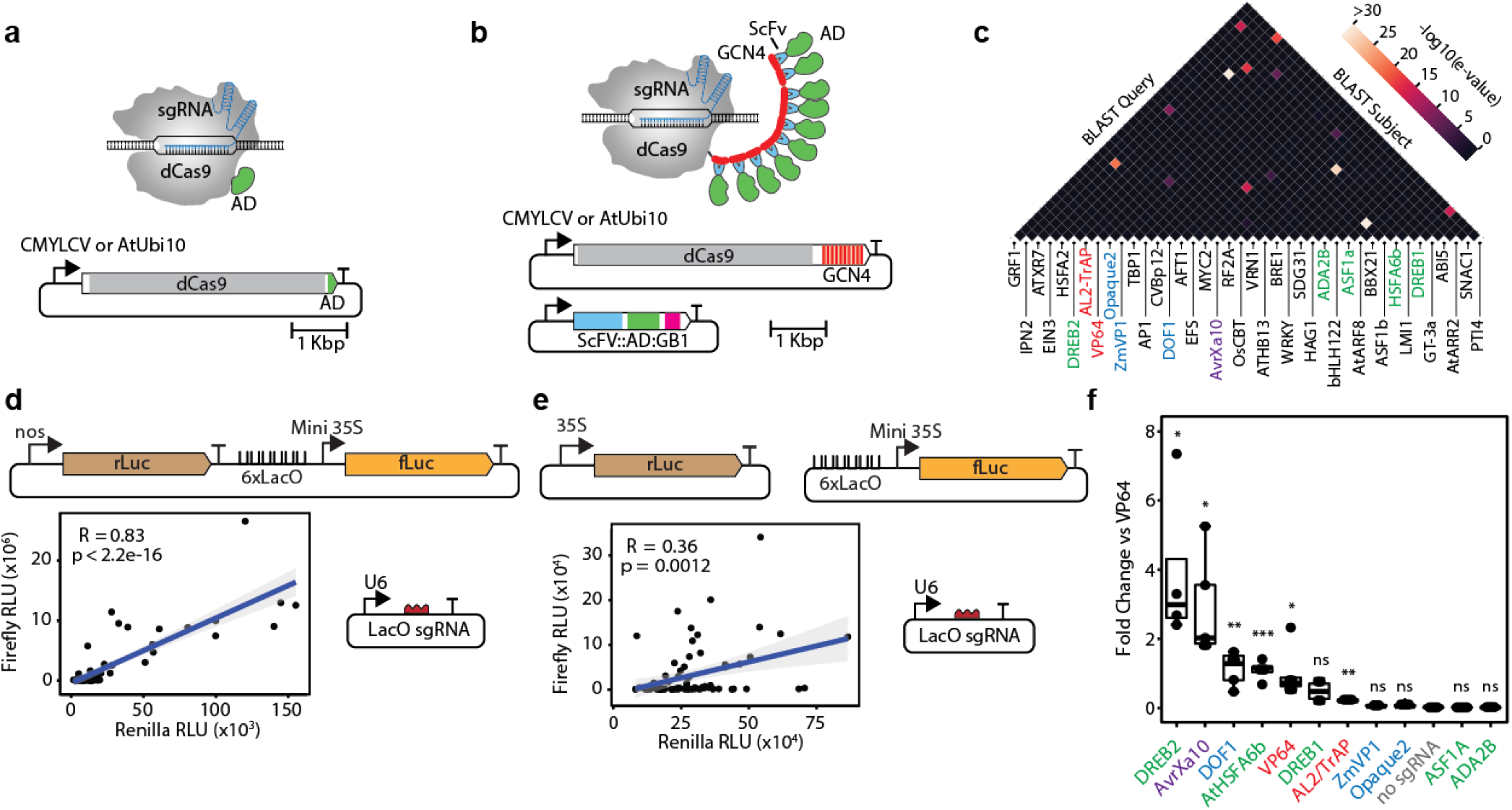
Screening a library of plant-derived activation domains with the dual luciferase protoplast assay. Cartoon images of dCas9-based Programmable Transcriptional Activators (PTAs) including the direct fusion (a) and SunTag scaffolding system (b). AD, activation domain; ScFv, single-chain variable fragment; GB1, solubility tag; GCN4, epitope bound by ScFv. (c) Heatmap showing the similarity of tested activation domains. Black denotes no significant similarity. Color-coded ADs highlight the strongest from the analysis in (f), with the following relationship: green, dicot; blue, monocot; purple, bacterial; red, viral. (d) Genetic construct design for a single-plasmid dual luciferase assay (top) and observed correlation between expression of PTA-controlled firefly luciferase and constitutively expressed Renilla luciferase (bottom). Data is from *S. viridis* protoplasts. (e) Genetic construct design for two-plasmid dual luciferase assay (top) and corresponding low correlation between firefly and Renilla luciferase expression (bottom). Data is from *A. thaliana* protoplasts transfected with the plasmids shown, plus the two plasmids encoding components of the SunTag system in (b). (f) A rank order of the strongest ADs is show for *A. thaliana* protoplasts against the VP64 positive control. Parametric Welch’s t-tests were performed comparing each AD with the negative control of scFv-VP64 delivered without a guide (no sgRNA). * corresponds to p <= 0.05, ** corresponds to p <= 0.01, *** corresponds to p <=0.001, ns is not significant (p > 0.05).

At one day post-transformation, all PTAs increased the expression of *WUS* compared to a no-sgRNA control. While dCas9-TV outperformed dCas9-VP64, the SunTag-VP64 design outperformed both direct-fusion PTAs by two orders of magnitude (Supplemental Figure 1). Given these results, we reasoned that testing a library of putative ADs as fusions to scFv in the SunTag system would result in a dynamic range of activation for AD comparison.

Next a list of ADs was compiled to test in plants. We started with a literature search of plant-derived transcription factors with known DNA binding domains from diverse transcription factor families. We computationally removed native DNA-binding domains, as these regions are highly conserved and are expected to produce off-target effects if retained in PTA fusions. If an activation domain was not empirically determined, we selected motifs enriched in acidic and/or aromatic residues. Acidic and aromatic residue patches are often associated with transcriptional activation domains due to their propensity to form phase separation condensates upon recruitment to a core promoter (24). We also added plant-evolved activation domain sequences from plant pathogen effector proteins such as TALE proteins from *Xanthomonas*, along with plant-derived sequences from transcription preinitiation complexes such as 14-3-3 scaffolding proteins. The terms “plant-derived” and “plant-evolved” are used here to refer to sequences identified in higher plants or plant pathogens, respectively.

The name, sequence, and citations for each domain tested can be found in Supplementary Table 1. We observe low amino acid sequence similarity across coding sequences from which ADs were derived, as illustrated by an all-by-all BLAST analysis (Figure 1c). This highlights the aim of this screen to survey a diverse array of plant-related ADs, versus optimizing on a well-performing family.

Putative AD sequences were codon optimized based on average codon usage tables for *A. thaliana* and *Oryza sativa* for reliable expression across a variety of plant backgrounds and synthesized as dsDNA fragments. We used Type IIS restriction enzyme cloning to assemble putative AD coding sequences into an scFv destination vector. ADs were expressed as C-terminal fusions to scFv, separated by a flexible Glycine-Serine (GS) linker containing Gly-Gly-Gly-Gly-Ser (25). AD coding sequences that failed multiple attempts to assemble in the scFv vector were removed from the study. ADs that failed multiple cloning attempts are denoted by an asterisk in Supplemental Table 1.

### A set of 5 plant-derived ADs show comparable activity to VP64 in a dual luciferase protoplast assay

A dual luciferase reporter assay was designed to screen the library for domains capable of activating transcription in plant cells (26, 27). On the luciferase reporter plasmid, we designed a synthetic promoter comprising six copies of the *Lac* operator (*Lac*O) upstream of the minimal *35S CaMV* core promoter upstream of firefly luciferase (28). A sgRNA targeting the *LacO* sequence will recruit the dCas9-based transcriptional activators to each of the six *LacO* motifs, such that firefly luciferase expression reports activation domain strength. To provide an internal control of plasmid copy number, a Renilla luciferase controlled by a constitutive promoter was inserted in the same plasmid, upstream of and oriented in the same direction as the firefly luciferase (Figure 1d).

Alternative activation domains were expressed as scFv-AD fusions driven by a *35S* promoter, allowing for strong expression in both monocot and dicot protoplasts. The dCas9-24xGCN plasmid was expressed by an *AtUBI*10 promoter in *A. thaliana* or the viral *CmYLCV* promoter in *S. viridis* (Figure 1b).

The library was first tested in *S. viridis* protoplasts. We isolated and transformed 100,000 cells with 2 micrograms of DNA in 96 well plates. Each transformation received dCas9-24xGCN, the *LacO* sgRNA, the single-plasmid dual luciferase plasmid with *35S* driving Renilla luciferase, and a given scFv-AD. Each scFv-AD was independently transformed twice per protoplast isolation. Following two *S*.*viridis* protoplast isolations spanning four independent transformations, a correlation was seen between Renilla and Firefly luciferase activity and was particularly pronounced in some batches (Figure 1d, Supplementary Figure 2,3). This could be due to strong PTAs at the 6x Lac operator acting as enhancers for the constitutive nos promoter on the vector. A two-plasmid dual-luciferase construct diminished this correlation (Figure 1e). With this experimental set-up, we compared the ability of alternative ADs to induce expression of Firefly luciferase. Across multiple experiments in different plant species, DREB2, DREB1, DOF1, AvrXa10, and HSFA6b showed good activity in the dual luciferase assay (Figure 1f, Supplementary Figures 4-6). Activity required the full SunTag system, as the ADs alone did not drive sub-stantial luciferase activity (Supplementary Figure 7). Activity of ADs was bench marked against scFv-VP64 delivered both with and without a guide RNA for positive and negative controls respectively (‘VP64’ and ‘no sgRNA’, Fig 1f).

### DREB2, DOF1, and AvrXa10 plant-derived activation domains outperform VP64 across endogenous loci

We next tested the efficiency of HSFA6b, AvrXa10, DOF1, DREB1, DREB2, and VP64 at endogenous loci. Single guide RNAs expressed from the *A. thaliana* U6-26 promoter were constructed to target the core promoters of *FLOWERING LOCUS T* (*FT*), *CLAVATA3* (*Clv3*), *PRODUCTION OF ANTHOCYANIN PIGMENT 1* (*PAP1*), *WUSCHEL* (*WUS*), and *LEAFY COTYLEDON1* (*LEC1*) for activation (Figure 2a-b, Supplemental Figure 8). These target genes were selected either because they are expected to drive a clear phenotype upon overexpression (*FT*), or because they are putative targets for the development of genetic biocontainment via Engineered Genetic Incompatibility, which is a longer term goal of our group (29). All sgRNAs used in this study are previously published (15, 30), except for the *WUS* guide, which was developed for this study. Approximately 500,000 *A. thaliana* protoplasts were isolated and transformed with the SunTag PTA and *A. thaliana* U6-sgRNA plasmids, with either two or three transoformation replicates per activation domain per target gene. After 24 hours, RNA was isolated and RT-qPCR was performed to quantify gene activation. We compared the activity of each AD against a no AD control for every sgRNA tested, in which the dCas9 and sgRNA were transformed without an scFv-AD. For each sample, the target gene and a control housekeeping gene (*PP2A*) were amplified. We report expression relative to the housekeeping gene *PP2A* in Figure 2b for *WUS, CLV3, FT*, and *PAP1*. By graphing all data points relative to the housekeeping gene, it is easier to observe both the fold-change in gene expression (height of the bar), and the absolute change in gene expression (location relative to the y-axis). This is important, as the fold-change in expression that can be achieved by PTAs is inversely correlated with the basal expression level. Genes already expressed at high levels cannot be overexpressed to the same relative degree as genes with a low basal expression level (10).

**Fig. 2.**
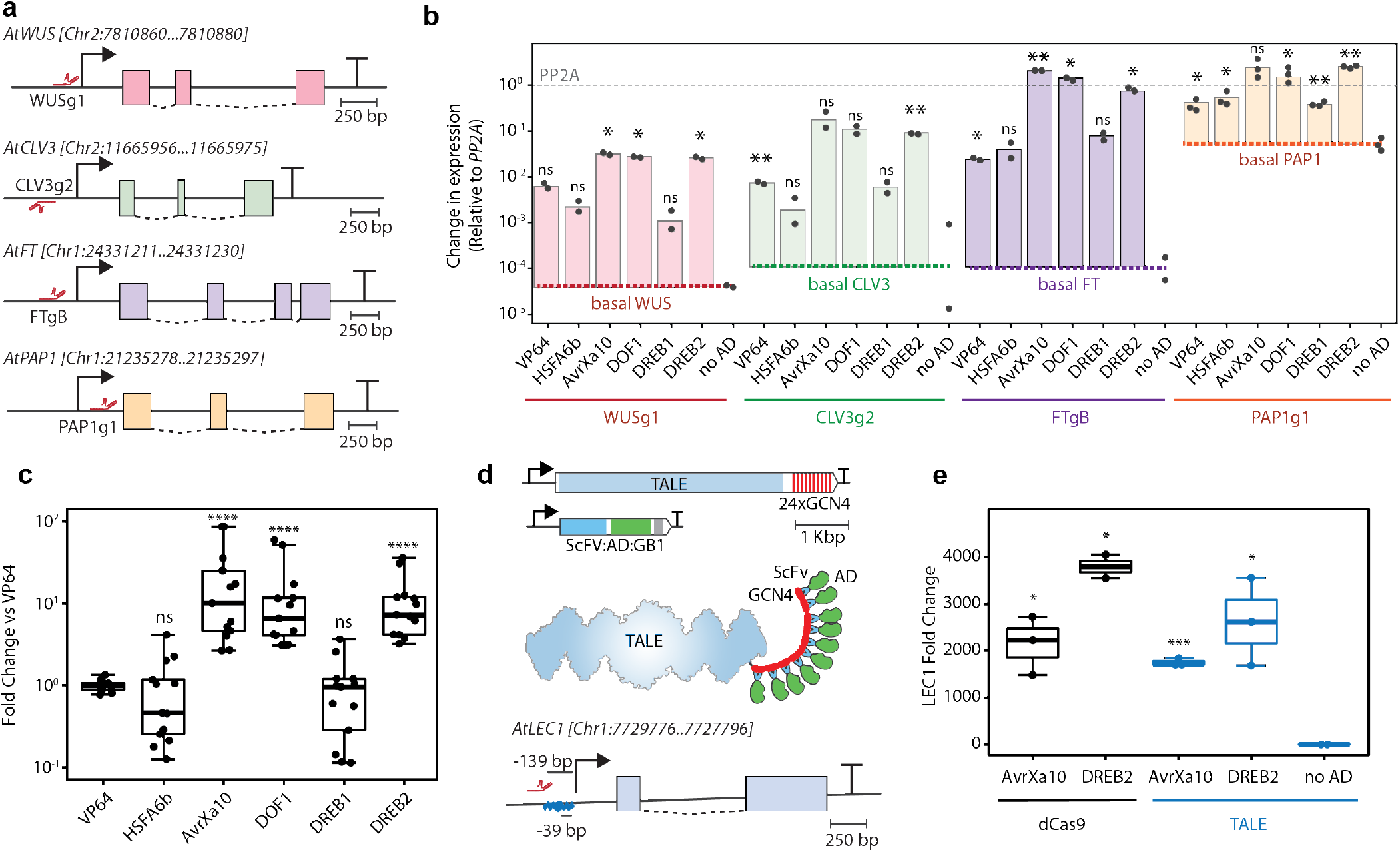
Endogenous gene activation in *A. thaliana* protoplasts. (a) Guide RNA design for a given promoter is shown for *WUS, CLV3, FT*, and *PAP1* loci. The precise location of sgRNA binding is given in brackets above each gene diagram. (b) Gene activation data for each corresponding gene is shown relative to the housekeeping gene *PP2A*. Basal level of gene expression, corresponding to values measured from no AD controls is unique for each target and is highlighted with colored, dashed line. Statistical significance compared to no AD control was determined conservatively using a Welch’s t-test due to low replicate numbers. See symbol key for p-values at the end of this legend. (c) Aggregate gene activation across all genes tested, where each dot represents an independent protoplast transformation. Parametric Welch’s t-tests were performed comparing each AD with the negative control NoAD. (d) The TALE DNA binding domain was fused to the GCN4 epitope tail to comprise a TALE-SunTag system (top). The TALE DNA binding domain was engineered to target the LEC1 promoter, compared to dCas9 binding the same promoter (bottom). The abbreviated names in this panel are described in the legend of Figure 1b.(e) The *LEC1* target gene in *A. thaliana* was targeted for gene activation comparing dCas9-SunTag (black) with TALE-SunTag (blue). * corresponds to p <= 0.05, ** corresponds to p <= 0.01, *** corresponds to p <=0.001, and **** corresponds to p<= 0.0001

The sgRNAs used in these experiments bind the core promoter regions, typically located 0-300 base pairs upstream of an annotated transcription start site (TSS) for a given gene except for the *PAP1* sgRNA used in this study which binds downstream of the annotated TSS. A clear pattern emerged across all sgRNAs tested, with AvrXa10, DOF1, and DREB2 consistently producing greater gene activation than VP64, HSFA6b, and DREB1 (Figure 2b, Supplemental Figure 8).

As expected, the maximum fold-overexpression was correlated with basal expression level. For example, *FT* saw a much greater fold-overexpression compared to no-AD control (>10,000) than did *PAP1* (50). However this is due to *FT* starting at much lower basal expression levels, and maximum absolute expression attained is comparable. *WUS*, on the other hand, could only be overexpressed to 10% the level of *PP2A*.

We compared each AD’s gene activation across the panel of endogenous genes to VP64. After setting VP64 activation values for each gene tested equal to 1, we combined data sets for all sgRNAs across four target genes (n=13). While the rank order changed depending on target gene, AvrXa10, DOF1, and DREB2 produced statistically significant gene activation across all target genes and guide RNAs that were tested when compared to VP64 (Figure 2c).

Because these activation domains described come from plant backgrounds, it is possible that transcriptional networks unrelated to the CRISPR-targeted promoter could be activated. To confirm the specificity of targeting endogenous genes in *A. thaliana* protoplasts, the transcriptional change of downstream target genes for each plant-derived AD were quantified in *A. thaliana* protoplasts after treatment with the dCas9 activation complex targeting the FT locus, and an unrelated target gene. These experiments were performed in *A. thaliana* protoplasts because HSFA6b, DREB1, and DREB2 are sequences taken from endogenous Arabidopsis genes. Downstream target genes for HSFA6b, DREB1, and DREB2 have been identified in the literature (31–36). Primers were designed to quantify the expression of these downstream target genes with RT-qPCR (Supplemental Table 5). The expression of a downstream target gene for the negative control (no AD) was compared to each AD treatment. No statistically significant off-target activation was detected for HSFA6b, DREB1, or DREB2 (Supplemental Figure 9). These results suggest specificity of the plant-derived ADs for the sgRNA-defined target site when fused to dCas9 in the scaffolding SunTag system.

We next tested the portability of these ADs between different programmable DNA binding domains. The 24xGCN4 epitope tail from the SunTag system was cloned as a C-terminal fusion to a TALE programmable DNA binding domain (Figure 2d). A TALE DNA binding domain is engineered by assembling a series of 20 TALE repeats, each of which contain a unique Repeat Variable Diresidue (RVDs) (15). Each RVD within a TALE repeat specifies a single nucleotide for binding, such that a specific 20 base pair genomic target sequence is specified by 20 RVDs. We built a TALE repeat targeting a 20 bp region 39 bp upstream of the transcription start site for the *A. thaliana* gene *LEC1*. In addition, we cloned a sgRNA targeting a 20 bp region 139 bp upstream of the transcription start site for the same *LEC1* target gene (Figure 2d). We then tested the ability of the AvrXa10 and DREB2 ADs to activate the *LEC1* target gene in conjunction with either dCas9 or TALE DNA binding domains in *A. thaliana*. As expected the dCas9 SunTag system using AvrXa10 and DREB2 domains resulted in strong *LEC1* gene expression with 2100 or 3800 fold greater activation than the No AD negative control (Figure 2e). The TALE SunTag system using AvrXa10 and DREB2 domains also resulted in strong *LEC1* gene expression of 1700 and 2600 fold greater activation than the No AD negative control. These results demonstrate the flexibility of the AvrXa10 and DREB2 ADs to drive strong expression of target genes using different programmable DNA binding domains.

### AvrXa10 activation domain promotes early flowing in transgenic plants

Next, we tested whether our plant-derived ADs would produce a phenotype upon gene activation in stable transgenic plants. Binary vectors were generated expressing the plant-derived ADs using a new PTA architecture named the MoonTag system (37). The MoonTag system utilizes a similar mechanism as SunTag, in which an epitope-nanobody interaction recruits the AD to the dCas9 molecule (38). In our Moontag system there are 24 copies of the GP41p epitope fused to dCas9, while AvrXa10, DOF1, and DREB2 were fused to the GP41 2H10 nanobody. Two sgRNAs targeting the *FT* core promoter in *A. thaliana* were expressed from *A. thaliana* U6 promoters to drive *FT* gene activation (Figure 3a-b). Plants were floral dipped and T1 transgenic lines were identified using the pFAST *Oleosin1* embryo oil body fluorescent reporter (39, 40). For comparison, we used a validated MoonTag-VP64 transgenic line as the positive control (37) and wild type Columbia-0 (Col-0) or sgRNA-only plants as the negative controls in these experiments (Figure 3c). T1 transgenic seeds were germinated on 1/2 MS agar plates and 1-5 seedlings from AvrXa10, DOF1, DREB2, and Col-0 genotypes were transplanted to soil. Four of 5 AvrXa10 plants displayed an early flowering phenotype, compared to a single early flowering plant of each DOF1 and DREB2 genotypes and none from the Col-0 controls (Supplemental Figure 10). We allowed these plants to set seed and used transgenic plants (without determining hetero- or homozygosity) from the T2 generation for molecular and phenotypic quantification of flowering time.

**Fig. 3.**
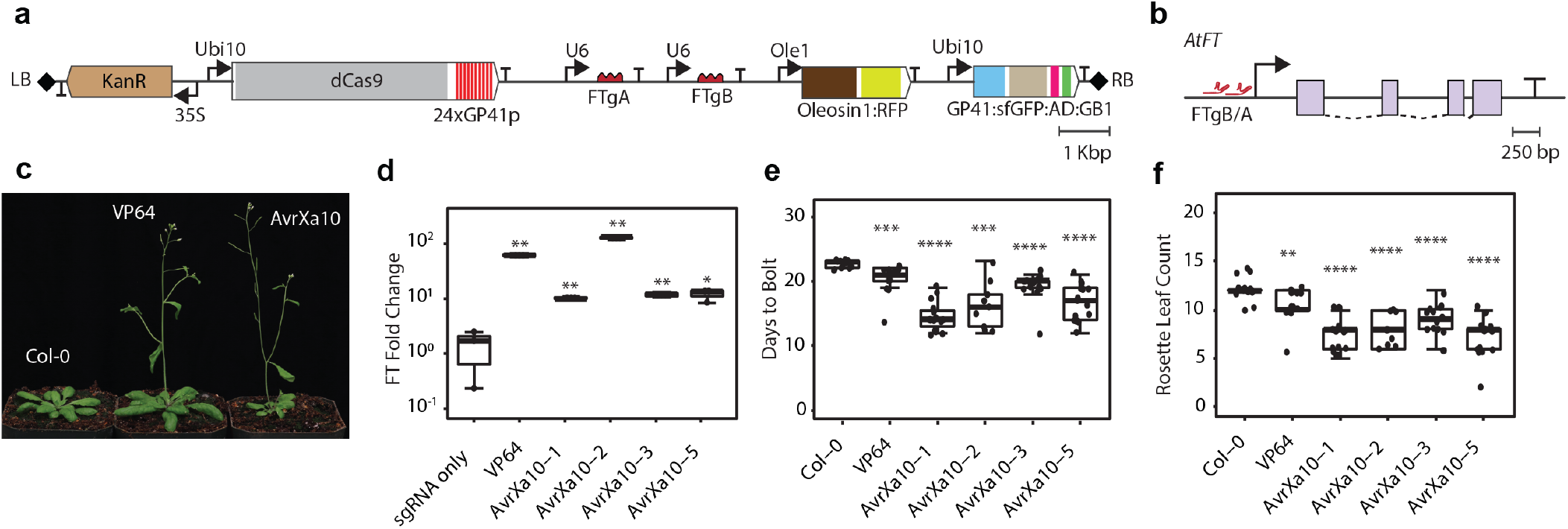
FT gene activation in transgenic plants. (a) Genetic construct design for binary vectors floral dipped to generate transgenic *A. thaliana* plants. The abbreviated names in this panel are described in the legend of Figure 1b, in addition to the following: RFP, red fluorescent protein; GP41, camelid nanobody; GP41p, peptide epitope for camelid nanobody; sfGFP, super-folder green fluorescent protein; LB and RB, left and right border, respectively. (b) Illustration of FTgA [Chr1:24331368..24331387] and FTgB [Chr1:24331212..24331230] design relative to the *FT* TSS. (c) Representative images of T2 transgenic plants for Col-0, VP64, and AvrXa10-1 transgenic lines. (d) Molecular quantification (RT-qPCR) of *FT* gene activation from 10d old T2 seedlings. Parametric Welch’s t-tests were performed on triplicate technical replicates from six pooled plants from each genotype compared to the sgRNA only negative control. (e) Bolting time quantification in T2 transgenic plants. (f) Rosette leaf quantification one day after bolting in T2 transgenic plants. Parametric Welch’s t-tests were performed in panels (e) and (f) comparing each line with the negative control Col-0. * corresponds to p <= 0.05, ** corresponds to p <= 0.01, *** corresponds to p <=0.001, and **** corresponds to p<= 0.0001

Sets of 20 seedlings per line were planted on selection media and allowed to germinate. Ten days after germination, seedlings were pooled into groups of six and RNA was extracted from the pooled group. We measured *FT* gene expression in the transgenic lines compared to a control line expressing only sgRNAs against a different target gene. The line used as a negative control expressed only sgRNAs, and lacked a Cas9 altogether. A binary vector map for this negative control line can be found in Supplemental Table S4. We observed strong gene activation in the positive control VP64 line, along with one of the AvrXa10 transgenic lines (Figure 3d).

We also quantified the *FT* overexpression phenotype in the T2 population by selecting 18 RFP-positive seedlings and planting them directly in soil for each transgenic line. Following germination, we waited for the seedlings to produce their first set of true leaves and used this time point as t=0. Two commonly used metrics of flowering time are bolting time and rosette leaf number at time of flowering (41, 42). We noted the day at which bolting was observed and calculated the difference between this day and t=0 to quantify bolting time (Figure 3e). We then counted rosette leaves one day post bolting for each seedling as an additional quantification of the *FT* phenotype (Figure 3f). We observed *FT* activation phenotypes across multiple AvrXa10 transgenic lines, with early bolting and reduced rosette leaf number in comparison to wild type plants. We also noted reduced rosette leaf size in several of the AvrXa10 plants (Figure 3c) compared to Col-0 and VP64 plants. We did not observe strong early flowering phenotypes for DOF1 and DREB2, likely due to a small number of lines recovered. Taken together we observed the plant-derived AvrXa10 AD functions and drives the strongest early flowering phenotype in *A. thaliana*.

### PTA-Mediated Gene Enhancer Activation

Finally, we demonstrated the ability of our improved plant PTAs to map enhancers. Co-targeting PTAs to both a core promoter and enhancer region has previously been shown to boost observed expression levels in human cells (43). To illustrate this concept in plants, we utilized the *FT* gene activation platform. Of the two sgRNAs tested, FTgB was capable of driving stronger gene activation than FTgA at the core promoter in our protoplast assays (Figure 2b, Supplemental Figure 8). According to previous literature, Block B (-1.8kb), Block C (-5.3kb), and Block E (+3.8kb) are three known enhancers of *FT* located either upstream or downstream of the TSS (44). We designed sgRNAs that target the centers of each 350-400 bp enhancer, along with the *FT* core promoter guide FTgB previously tested. In addition, we designed guides that target sequences located between the annotated enhancer regions and the core promoter (Figure 4a).

**Fig. 4.**
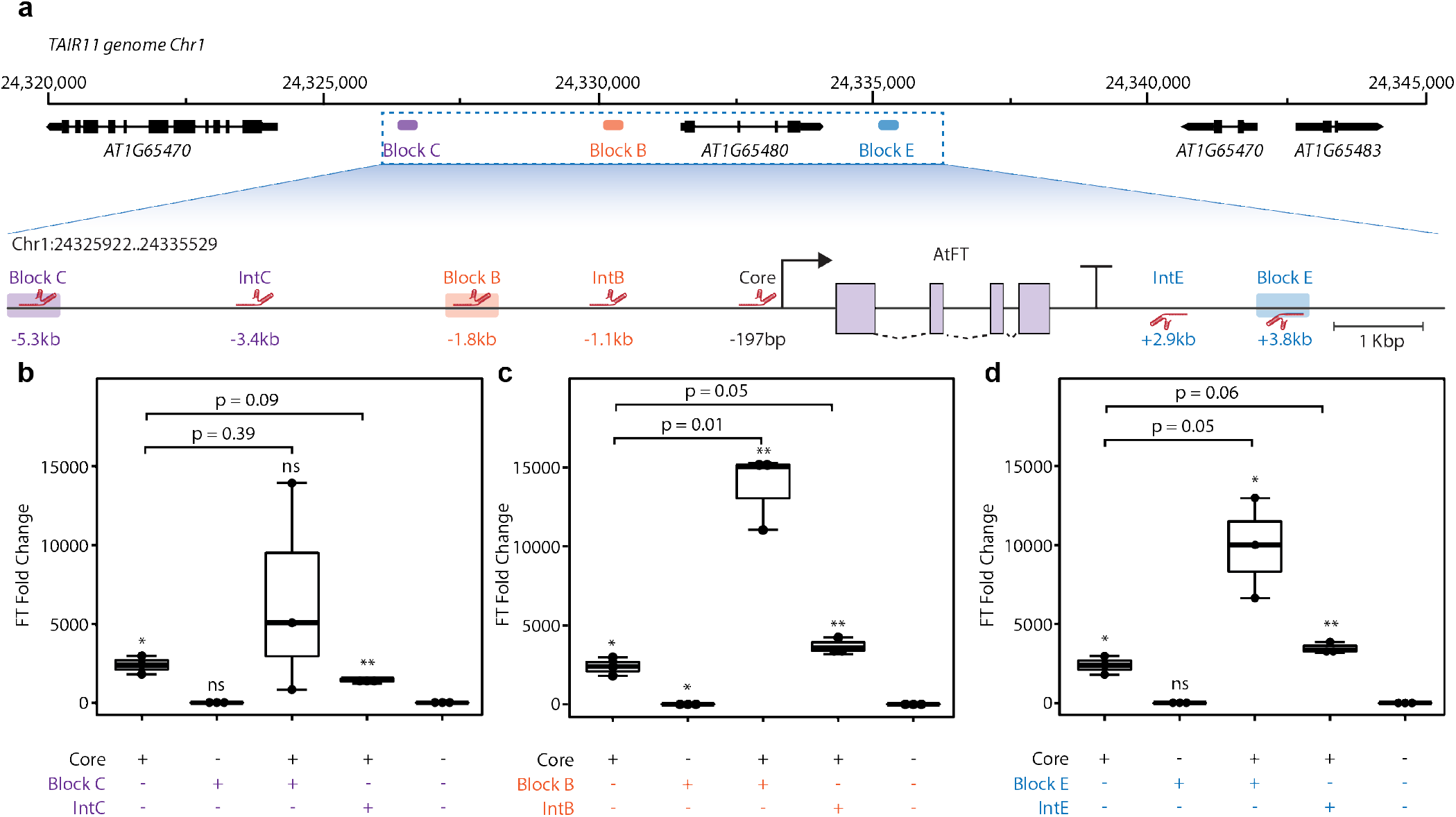
PTA-mediated enhancer activation at the *A. thaliana FT* locus using DREB2. (a) Schematic illustration of sgRNA design in relation to the region of *A. thaliana* chromosome 1 surrounding the *FT* gene. Top depicts entire 25kb locus and bottom shows more detail of enhancer location and sgRNA design. sgRNAs drawn above or below the line denote targeting the sense or antisense strand, respectively. Distances labeled are relative to the *FT* TSS. (b-d) *FT* gene activation, measured by RT-qPCR when targeting Block C (b), Block B (c) or Block E (d) with different combinations of sgRNAs (x-axis). Data plotted in b-d are all from a single experiment but have been organized into three separate subplots (with the same data from the core promoter and negative control shown in each subplot) to facilitate interpretation of the results. Parametric Welch’s t-tests were performed comparing each sgRNA combination with the negative control where no sgRNA is delivered. P-values for these analyses are denoted symbolically, where * corresponds to p <= 0.05, ** corresponds to p <= 0.01, and ns means ‘not significant’ (p > 0.05). Brackets with specific p-values listed are separate parametric Welch’s t-tests comparing the groups below the bracket ends.

For each of Blocks B, C, and E, targeting the enhancer alone did not result in overexpression above no-sgRNA controls (Figure 4b-d). Targeting both the core promoter and the enhancer led to a significant increase in expression level for Blocks B and E. The Block C enhancer-targeting sgRNA produced greater expression levels in two of three replicates, but a large amount of variance in this experiment prevented it from reaching statistical significance. When targeting enhancers plus core promoters, mean fold overexpression compared to no-sgRNA controls ranged from 6,000-fold (Block C) to 14,000 fold (Block B). This improvement over targeting the core promoter alone (2,400-fold overexpression compared to control) could not be attributed to having a second sgRNA, as the sgRNAs targeting sequences outside of the enhancer regions did not significantly boost expression levels (Figure 4b-d).

Finally, we tested the specificity of PTA-mediated enhancer activation by quantifying the expression of the FAS1 gene (AT1G65470) located upstream of the FT gene (AT1G65480). It is possible that the Block B, C, and E regions of open chromatin serving as enhancers for the FT locus may also act as enhancers for nearby genes when targeted with dCas9-based activators. To test this, RT-qPCR primers were designed for the FAS1 transcript, and its expression was measured across all enhancer treatments. The enhancer treatments surveyed for FAS1 expression in the study were targeting of the FT core promoter alone, co-targeting of the FT core promoter + Block B, C, and E, and finally the FT core promoter sgRNA + dCas9-24xGCN4 lacking any AD. Little activation was observed at the FAS1 gene when targeting the FT core promoter along with guides targeting the open chromatin enhancer regions of Block B, C, and E when compared to the No AD negative control (Supplemental Figure 11).

## Discussion

We set out to identify plant-derived ADs that function as well or better than VP64 in plant systems. Through a library approach we were able to clone and test putative ADs in protoplast dual luciferase assays to determine which sequences showed activity comparable to VP64. We identified a core set of 5 activation domains comprised of AvrXa10, HSFA6b, DREB1, DREB2, and DOF1 that showed strong activity in both the monocot model *S. viridis* and the dicot model *A. thaliana* dual luciferase protoplast assays. The high throughput nature of protoplast transformation serves as a valuable tool for testing large numbers of genetic parts (22).

While performing these assays, we observed what appeared to be local off-target activation of our dual luciferase reporter. DREB2, DREB1, and HSFA6b consistently produced the largest Renilla luciferase values, even though they were targeted to the firefly luciferase promoter. This resulted in a linear correlation between firefly luciferase and Renilla luciferase values (Figure 1d). Because this correlation was reduced by splitting the reporters into two plasmids (Figure 1e), we conclude expression from the upstream Renilla promoter is the result of trans-activation from the PTA. In this case, the promoter driving the Renilla gene was located approximately 3kb upstream of the sgRNA binding site. Interestingly, targeting a PTA to within 3kb of the endogenous *FT* locus did not cause *FT* overexpression unless at least one sgRNA bound to the core promoter region. It is possible that the proximity effect seen in our single-plasmid dual luciferase assay is dependent on transcriptional activity from the targeted promoter. Nucleosome formation on transiently-transfected, episomal DNA is known to be altered compared to chromosomal DNA(45–47), and this could allow for interactions between reporter construct promoters in our protoplast assay. This phenomena may vary depending on the locus, the number of sgRNAs, or even the AD that is directed to a target site. In genome-wide screens, off-target activation events are rarely seen (48).

We demonstrate endogenous gene activation in which AvrXa10, DOF1, and DREB2 consistently produce the greatest activation across all genes tested. It is interesting to note that the relative strength of ADs did vary both across species and across target loci. For example the AvrXa10 domain shows the largest range of activation among the three strongest ADs across target genes in *A. thaliana* protoplasts. Given that AvrXa10 is derived from a plant pathogen, perhaps there are varying degrees of host response to the expression of this domain across plant species and/or target gene.

We demonstrate the ability of AvrXa10, DOF1, and DREB2 plant-derived activation domains to drive gene activation of the *FT* locus in transgenic plants. T1 plants showed an early flowering phenotype, and this phenotype was maintained in the T2 generation where early flowering was quantified. This work illustrates the feasibility of moving from a protoplast-based discovery platform to validating the tools in stable transgenic lines. Changes in leaf size were also observed, particularly in the AvrXa10 expressing transgenic lines. Decreases in leaf size and number have been observed by others, deemed the “extra-early flowering” phenotype (49).

Finally, we showcase the ability of the DREB2 activation domain to function in a potential gene enhancer discovery pipeline. The *FT* locus contains three known enhancers of gene expression Block B, C, and E. These enhancers are located vast distances from the core promoter, and fail to produce any observable activation when targeted with dCas9 PTAs. While we do observe slight boosts in *FT* activation when co-targeting both the core promoter and an intervening region between enhancers, these boosts were rarely statistically significant and never as dramatic as the overexpression seen when co-targeting an enhancer. Noisy gene activation at distances outside of the core promoter (50-400bp upstream TSS) has been reported before (48), and may be a natural consequence of transcription pre-initiation dynamics.

Determining the location and activity of enhancer sequences can be challenging. Luciferase assays in protoplasts have traditionally been used to confirm enhancer activity by cloning an enhancer upstream of the firefly luciferase reporter and checking for activity (50). PTA-mediated enhancer activation retains the original genomic architecture and chromatin state of the enhancer-promoter pair, while traditional luciferase assays deploy enhancer sequences in non-native genomic architectures on plasmids which lack nucleosomes. Given our understanding of enhancers functioning through a DNA looping mechanism, retaining this genetic architecture may be important.

Once a putative enhancer is identified it can also be difficult to determine the promoter, or promoters, it regulates. Recent studies have utilized methods such as STARR-seq to confirm the transcriptional regulatory capacity of distal accessible chromatin elements in protoplasts (51). We anticipate PTA-mediated enhancer activation can be similarly used for testing distal accessible chromatin activity in plants. One could design a series of sgRNAs to target intergenic regions in tandem with a sgRNA targeting the core promoter for a gene of interest. Conversely, an accessible chromatin region of interest can be targeted along with the core promoters of nearby genes to identify which transcripts are significantly activated by the enhancer. When conducted in a high-throughput transient platform such as protoplasts, this pipeline could reveal novel enhancer-promoter interactions. This type of enhancer mapping can further expand the definition of gene by including enhancer regions in addition to the core promoter, untranslated regions, exons, introns, and terminator comprising a transcriptional block. Determining the general reliability of this method for mapping enhancers will require follow-up experiments in different genes.

This study has limitations that should be addressed with future work. The PTAs we describe here work great in research and development environments, but they have components derived from human or plant pathogens that would trigger special regulatory approval considerations if used in transgenic plants intended for field release. Replacing the VP64 domain with plant-evolved ADs mitigates that in part, however the dCas9, GB1, and AvrXa10 components are similarly derived from human or plant pathogens (52).

In addition, it has been noted that scFv proteins have difficulty forming disulfide bonds within the reducing environment of the cytosol, leading to poor solubility (53). For these reasons, replacing GB1 with alternate solubility tags could be addressed to further improve the performance of these PTAs in plants. For example, a naturally occurring solubility tag from spider silk shows promise as an alternative to the Streptococcal GB1 sequence (54). Improving the solubility of the PTA complex without compromising its transcriptional activation potency should be of significant interest moving forward.

With this work we showcase potent PTAs in plants capable of driving strong expression of target genes. We present three transcriptional activation domains from plant backgrounds that outperform VP64 in both reporter gene assays and targeting endogenous loci. We further showcase the portability of both DREB2 and DOF1 in diverse plants. They provide stronger expression than VP64 for target genes in *A. thaliana, S. viridis*, and *Z. mays*. Finally, we show that the DREB2 domain is equally as potent when fused to an alternative TALE programmable DNA binding domain. Given these results, we anticipate that these domains will be used by plant scientists to effectively activate transcription of target genes within diverse plant species.

## ACKNOWLEDGEMENTS

MHZ and DV are supported by Department of Energy Grant DE-SC0018277. MHZ, JACM, AS, and MJS are supported by the USDA grant 2018-33522-28747 and are supported by the Advanced Plant Technologies program, DARPA Award HR001118C0146. MHZ and AS are supported by an NIH NIGMS Biotechnology Training grant NIHT32GM008347. SCH was supported by a Doctoral Dissertation Fellowship from the University of Minnesota Graduate School.

## AUTHOR CONTRIBUTIONS

MHZ, DV, and MJS conceived this study. MHZ designed and built constructs, planned experiments, collected and analyzed data. ACM, AS, and MHZ performed protoplast isolation, transformation, RNA extraction, and RT-qPCR analysis. ACM built constructs and carried out floral dip. ACM provided VP64 transgenic line for *FT* analysis. MHZ, MJS, and DV wrote the manuscript.

## Supplementary Note 1: Materials and Methods

### Plasmid construction

Putative activation domains were selected from literature analysis for the potential to recruit transcription initiation machinery in plant cells. DNA binding domains from transcription factors were identified and removed on the basis of sequence conservation to experimentally determined DNA binding domain motifs. Patches of acidic/aromatic residues were identified as core motifs for a prospective activation domain, and a given sequence was selected to encompass as many of these patches as possible without including the DNA binding domain. Synthetic DNA fragments were then designed and codon optimized based on average codon usage in *A. thaliana* and *Oryza sativa*. Fragments were synthesized by Genscript with BsmBI cloning sites flanking the prospective activation domain. Fragments were cloned into an scFv destination vector with BsaI and BsmBI cloning sites generating compatible overhangs. The assembled plasmids generate an in-frame C terminal fusion of the activation domain to the scFv. The scFv is driven by the *CaMV 35S* promoter, and also contains a C terminal fusion of the GB1 solubility tag. The scFv-AD is recruited to targets by the dCas9-GCN4. This vector was generated by fusing 24 copies of the GCN4 epitope tag to the C-terminus of *A. thaliana* codon optimized dCas9. Each GCN4 repeat was separated by short linker GS linker sequences. The dCas9 is expressed under the AtUbi10 promoter in dicots, the *CMYLCV* promoter in *S. viridis*, or the *ZmUbi10* promoter in *Z. mays*. Guide RNAs were designed and expressed from *U6* promoters, either the *A. thaliana* U6 promoter for dicots or the *O. sativa U6* promoter for monocots. A complete list of plasmids used in this study can be found in Supplemental Table 1.

### PTA protein sequence similarity

BLAST (ver. 2.8.1+)(55) was used to perform an all-by-all comparison of the proteins from which the activators were derived. Full length amino acid sequences were used as both query and subject, and no e-value threshold was set. Expectation values were negative log transformed for plotting.

### Protoplast isolation and transformation

*S. viridis* and *A. thaliana* protoplasts were isolated and transformed as stated in previous protocols(21, 22). ME034 *S. viridis* plants are grown in a growth chamber set to 31/22 degrees C with 12h diurnal cycle. Mesophyll cells can be isolated from leaf tissue 14-21 days post germination. Col-0 *A. thaliana* plants are grown in a growth chamber set to 22C with 16h light cycle. Mesophyll cells can be isolated from leaf tissue 14-21 days post germination. *Z. mays* protoplasts were isolated and transformed as stated in previous protocols. (56) *Z. mays* plants were grown in a growth chamber at 25 degrees C under 16h light cycle. Prior to isolation, plants were ‘greened’ by first placing the seedlings in the dark for 5 days post-germination followed by 2 days of exposure to light prior to isolation. Plasmids to be transformed in all systems are prepared according to the manufacturer’s protocol (Qiagen 12945) to ensure transformation grade endotoxin free DNA.

### Dual luciferase assay

A dual luciferase assay was designed with the firefly and Renilla luciferase reporter genes from Promega (26). A Renilla luciferase was cloned upstream of a firefly luciferase on a single plasmid. The Renilla CDS was under the control of a constitutive promoter, either the *CaMV 35S* or weaker *Nos* promoter. Upstream of the firefly luciferase CDS was a minimal *CaMV 35S* promoter, containing 60 bp upstream of the TSS. Six copies of the *lac* operator were cloned upstream of the minimal promoter driving firefly luciferase. A sgRNA targeting the *LacO* region results in targeting of the dCas9 activation to the firefly luciferase promoter. We drove dCas9-24xGCN4 with a constitutive promoter (*CMYLCV, AtUbi10, or ZmUbi1*), the *LacO* sgRNA driven by *O. sativa U6* promoter, and the scFv with a constitutive *35S* promoter. We co-transformed *S. viridis, A. thaliana*, or *Z. mays* protoplasts with 20 micrograms each of the dCas9-24xGCN4, *Lac*O sgRNA, scFv-AD, and dual luciferase reporter plasmids. Following 24h incubation at 25C, we lysed the cells in 1X Passive Lysis Buffer from Promega. We then quantified firefly and Renilla luciferase luminescence using the GloMax Explorer plate reader (Promega GM3500) equipped with dual injectors with the Dual Luciferase Assay kit (Promega E1960). Firefly luciferase substrate was injected first and luminescence was quantified, followed by injection of the Renilla luciferase substrate and subsequent quantification. We normalized for delivery first by dividing firefly luciferase activity by Renilla luciferase activity unless otherwise described, followed by Fold Change calculation by dividing a given AD by the negative control or VP64. A complete list of plasmids used in this study can be found in Supplemental Table 2.

### RNA isolation and quantification

We isolate RNA from either protoplasts or leaf tissue according the the TRIzol manufacturer’s protocol (Thermo 15596026). For protoplasts we spun the cells down at 500g for 2 minutes, followed by removal of W5 buffer before re-suspending cells in 1mL of TRIzol. RNA was collected 24h post-transformation for protoplast samples. For plant tissue samples, we first snap froze the sample in liquid nitrogen followed by shaking in a paint shaker apparatus with metal beads to homogenize frozen tissue. We then add 1mL of TRIzol to the homogenized tissue. We then follow the TRIzol protocol according to manufacturer specifications. We then treat the samples with Turbo DNA Free Kit (Invitrogen AM1907) to remove any plasmid and/or genomic DNA from the RNA sample. To quantify a transcript we then perform RT-qPCR using gene-specific primer pairs. A list of primers used for quantification can be found in Supplemental Table 2. We follow a RT-qPCR cycling protocol as defined by the NEB Luna One-Step RT-qPCR kit manufacturer’s protocol (NEB E3005L). Primers are designed to have Tm values of 60C and yield amplicon lengths between 75-175bp. The primers typically span an intron, such that the shorter PCR product corresponds to spliced mRNA. We quantify gene expression for relative comparison between treatments using the delta delta Ct method. All primers can be found in Supplemental Table 2.

### Plant transformation and genotyping

To generate transgenic *A. thaliana* plants we performed floral dip protocols as previously described (39). Binary vectors were constructed containing an antibiotic resistance gene along with the pFAST Oleosin-RFP transgene (40). *A. thaliana* ecotype Columbia-0 was grown to maturity and floral dipped with Agrobacterium strain GV3101 transformed with the described binary vectors. The antibiotic selection used in this study was either Kanamycin or Bialaphos resistance. T0 seeds were identified by the pFAST system in which RFP-positive seedlings contain the transgene. We placed RFP+ seedlings directly onto soil, or onto selective media, to grow T1 plants. T1 plants were grown to maturity and allowed to set seed. All molecular characterization was performed on T2 lines, along with phenotyping. To quantify *FT* overexpression phenotypes in T2 plants, we planted 18 RFP positive T2 plants from each T1 parent. We chose time point 0 to be the date at which the first set of true leaves emerge from a seedling. We then count the number of days elapsed until bolting is observed. We also count the number of rosette leaves one day after bolting was observed. RT-qPCR was performed to quantify target gene expression, using primers indicated in Supplemental Table 2.

## Supplementary Note 2

**Supplementary Table S1.**
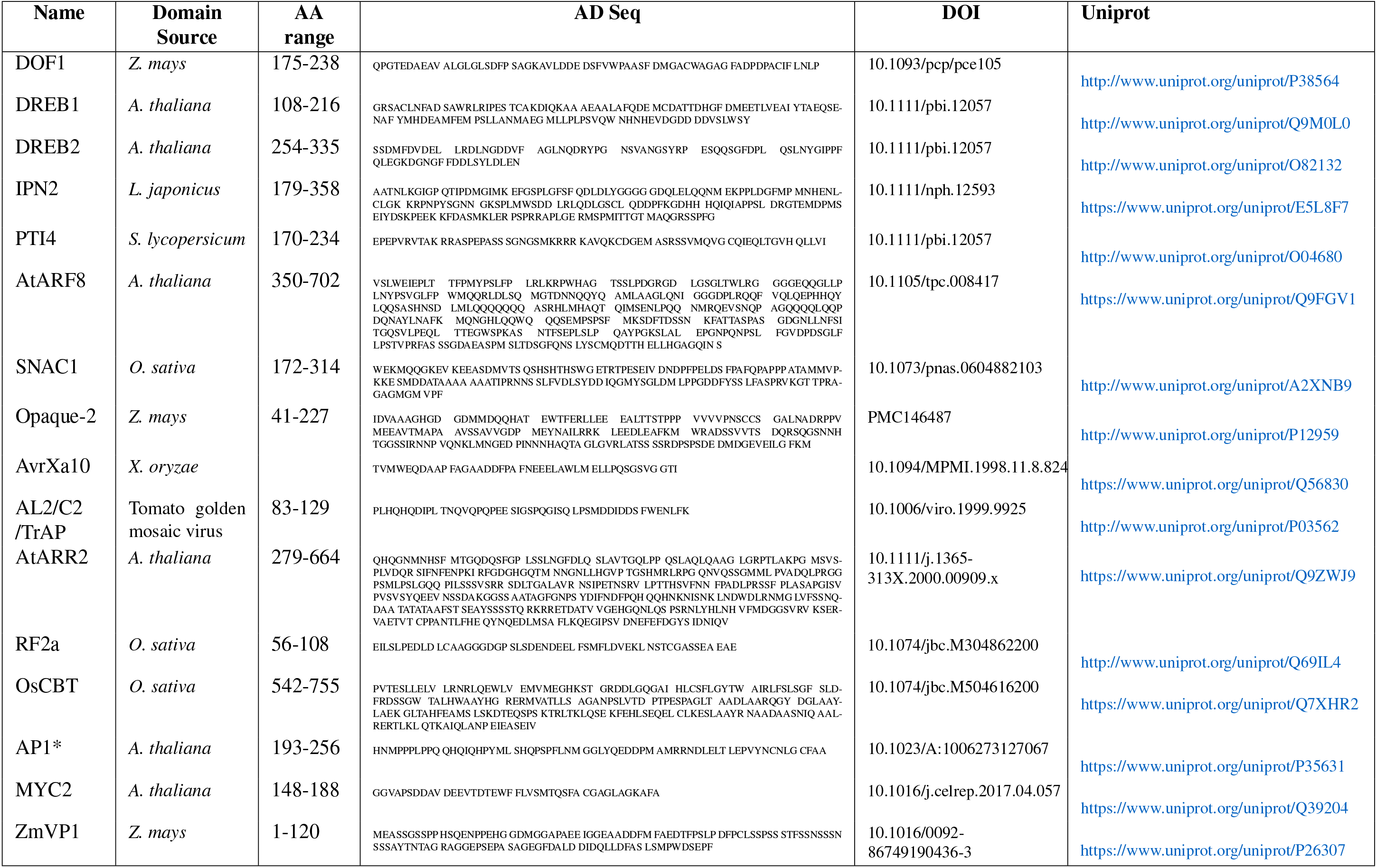
Activation domains.

## Supplementary Note 3

**Supplementary Table S1.**
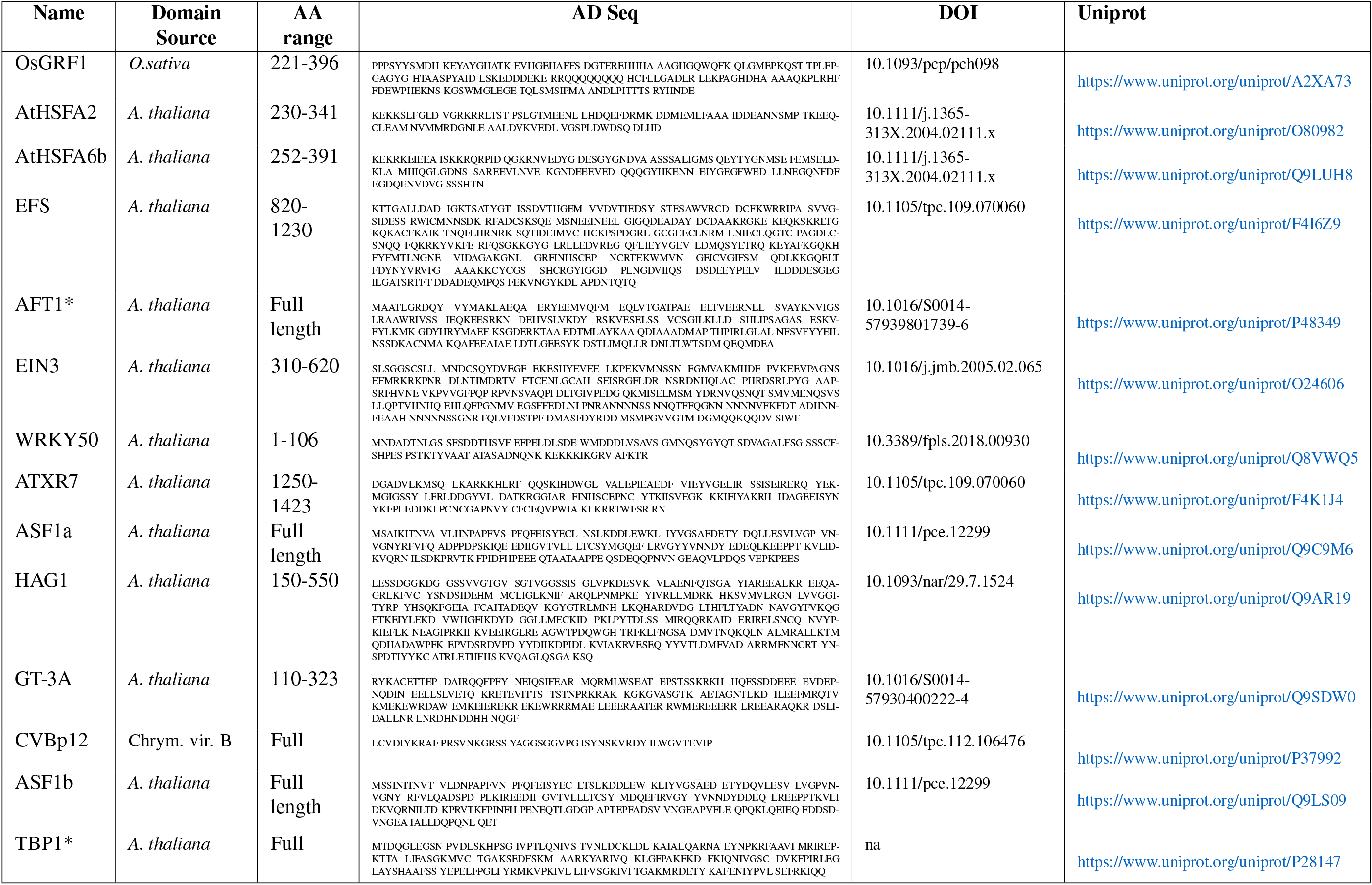
Activation domains continued.

## Supplementary Note 4

**Supplementary Table S1.**
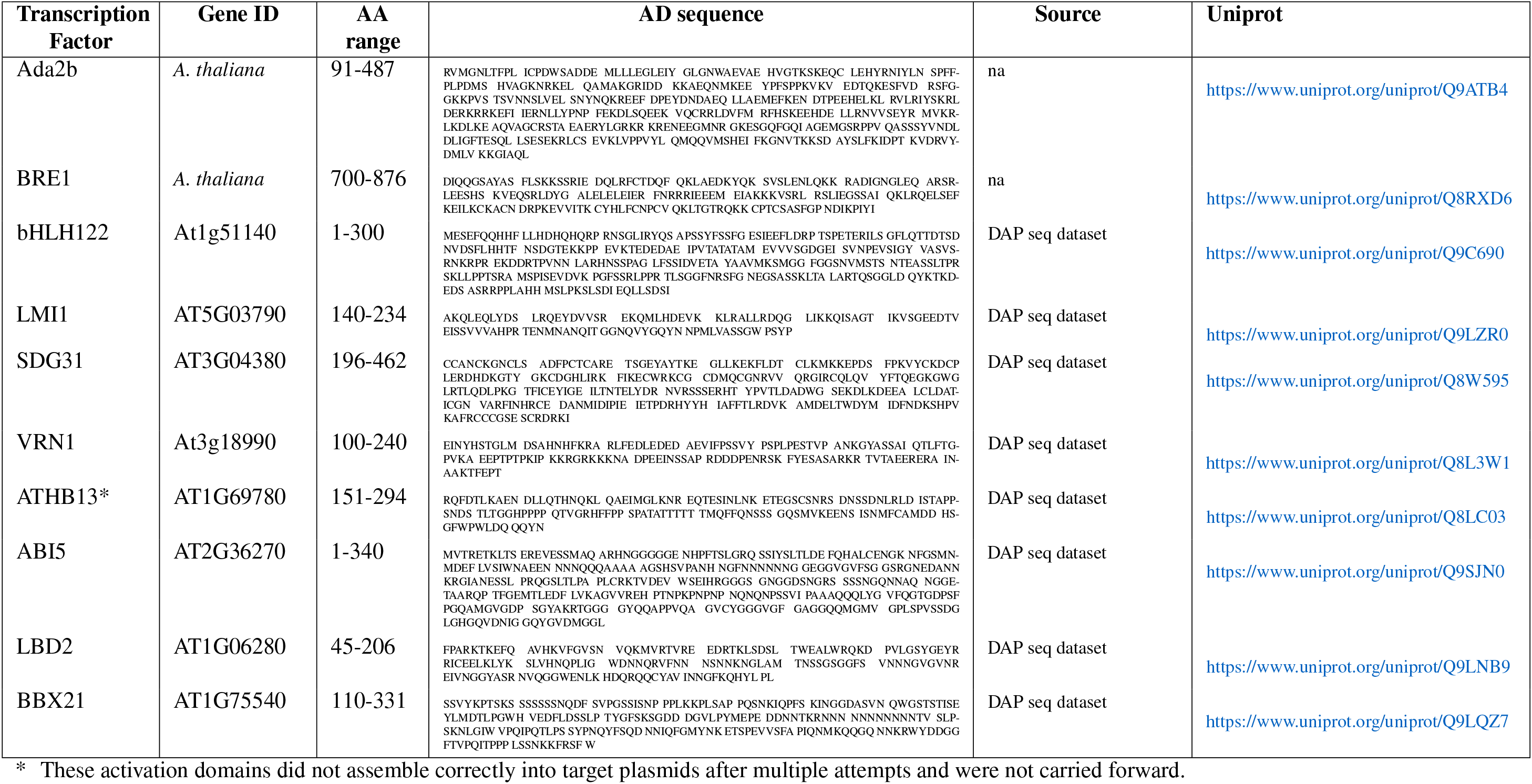
Activation domains continued.

## Supplementary Note 5

**Supplementary Table S2.**
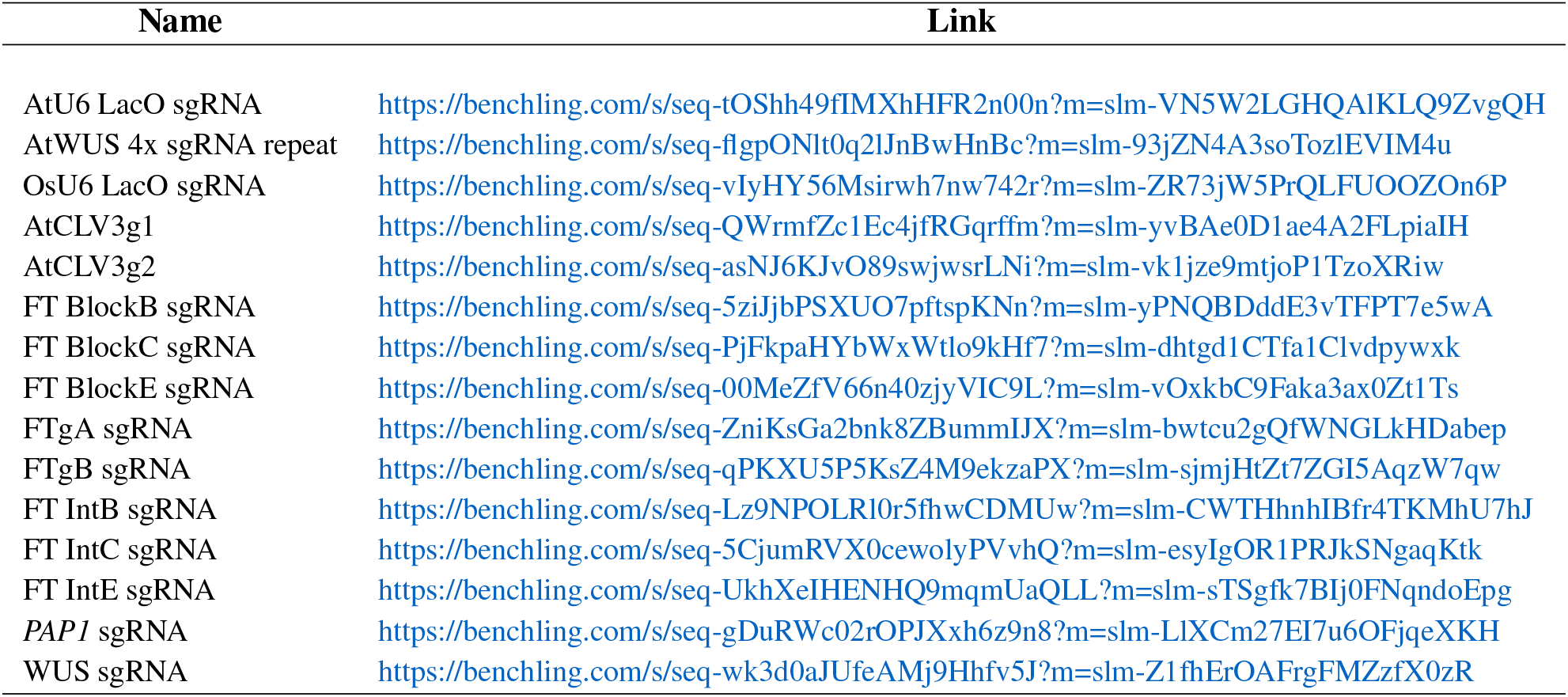
sgRNA constructs.

## Supplementary Note 6

**Supplementary Table S3.**
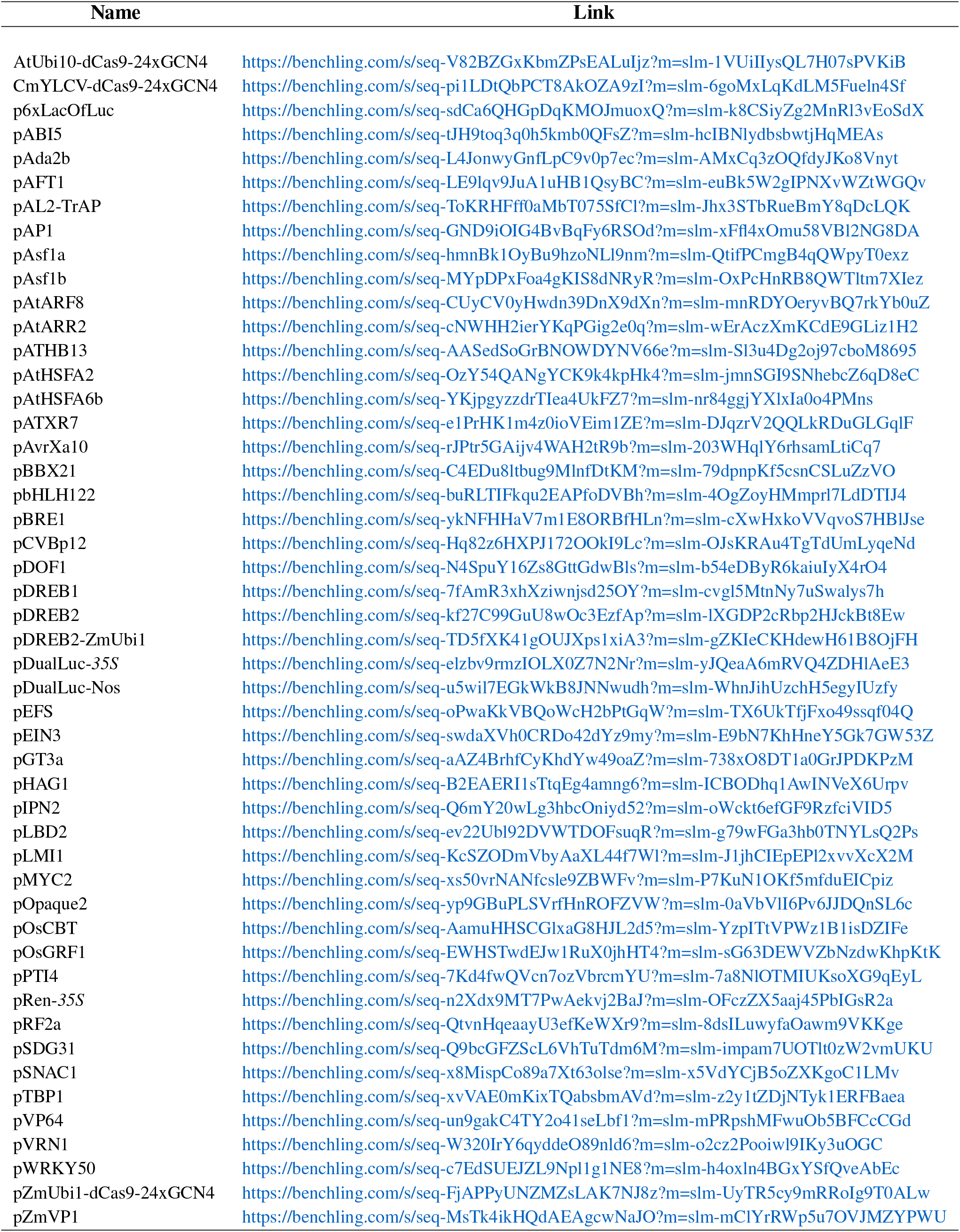
coding sequences.

## Supplementary Note 7

**Supplementary Table S4.**
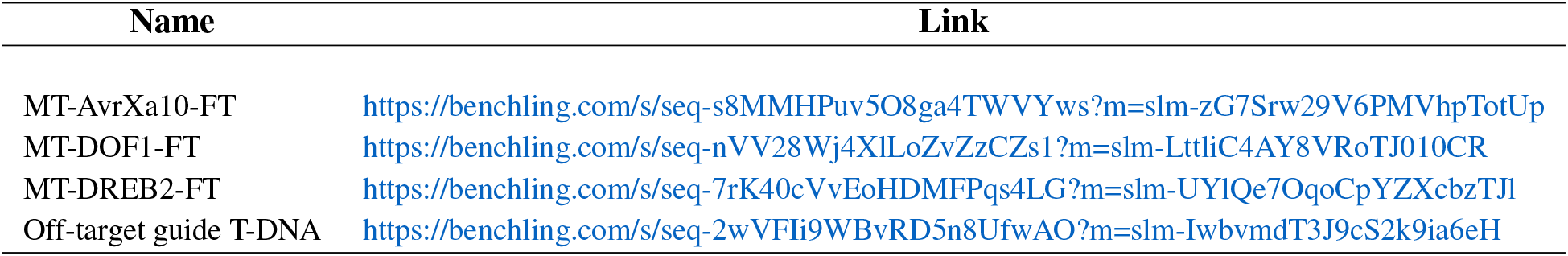
Binary vector maps.

## Supplementary Note 8

**Supplementary Table S5.**
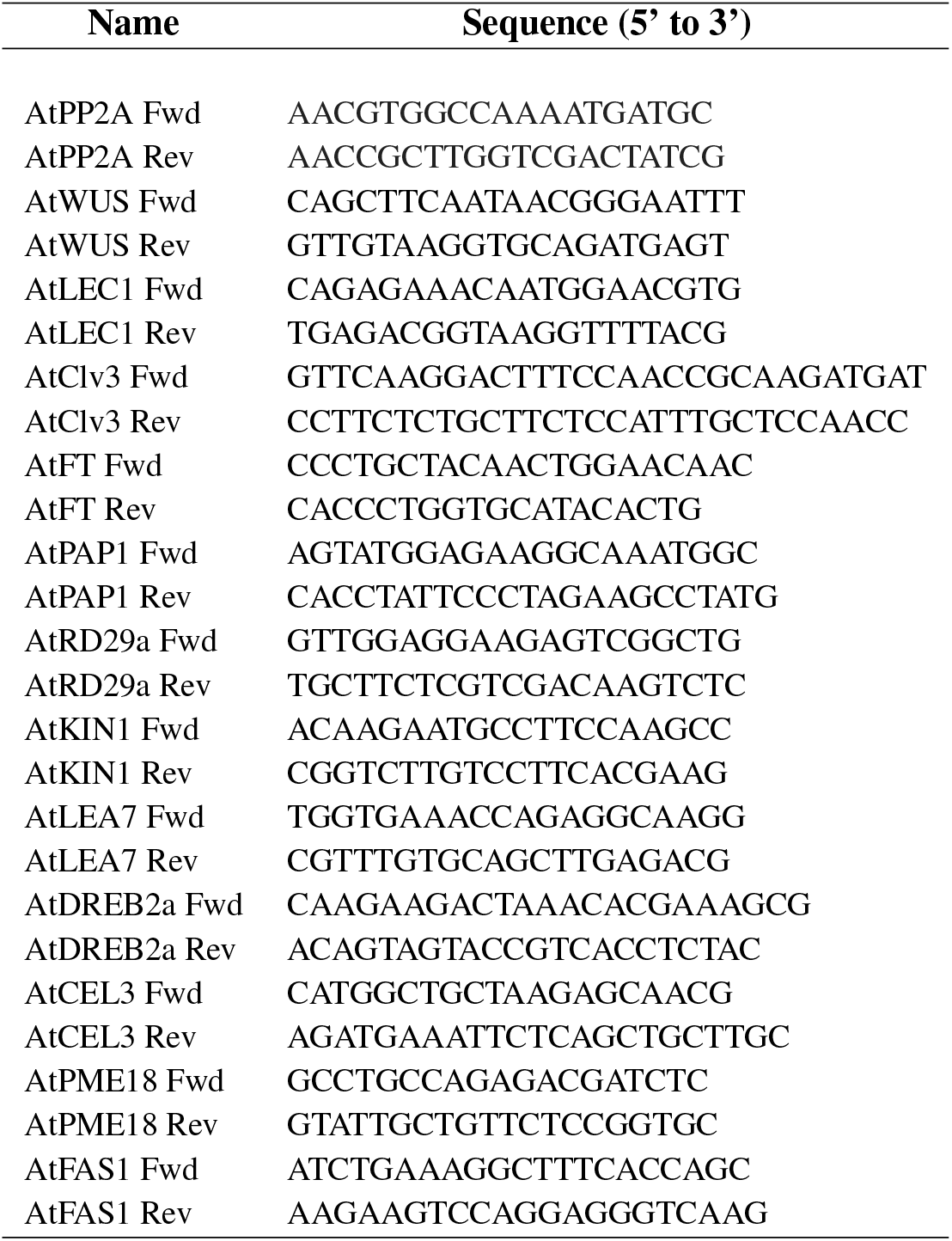
RT-qPCR primers.

## Supplementary Note 9: Supplementary Figure 1

**Supplemental Figure 1.**
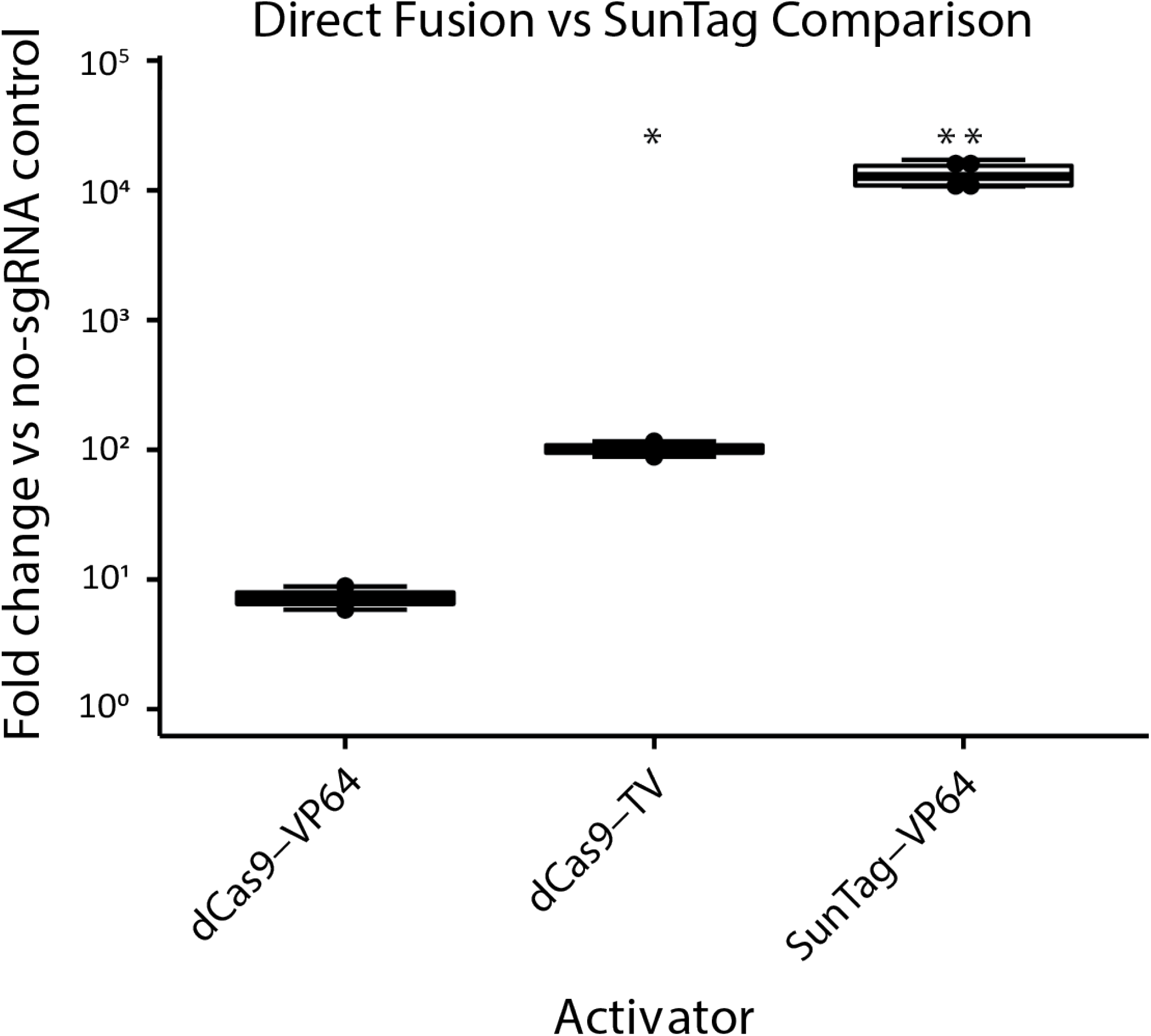
Direct fusion PTA vs SunTag PTA comparison in *A. thaliana* protoplasts. The direct fusion class of PTAs containing dCas9-VP64 and dCas9-TV, along with the SunTag-VP64 activator, was transformed into protoplasts with a 4x *Wuschel* sgRNA array targeting the core promoter for gene activation. RNA was extracted and activation was quantified by RT-qPCR using the delta delta Ct method to generate the box-and-whisker plots. Parametric Welch’s t-tests were performed comparing dCas9-TV or SunTag-VP64 with the dCas9-VP64 direct fusion at the same locus. A link to the sequence file of the 4X sgRNA repeat is given in Supplementary Table S2. All data is normalized against Ct values obtained using a no-sgRNA control (not shown). * corresponds to p <= 0.05, ** corresponds to p <= 0.01, *** corresponds to p <=0.001, and **** corresponds to p<= 0.0001.

## Supplementary Note 10: Supplementary Figure 2

**Supplemental Figure 2.**
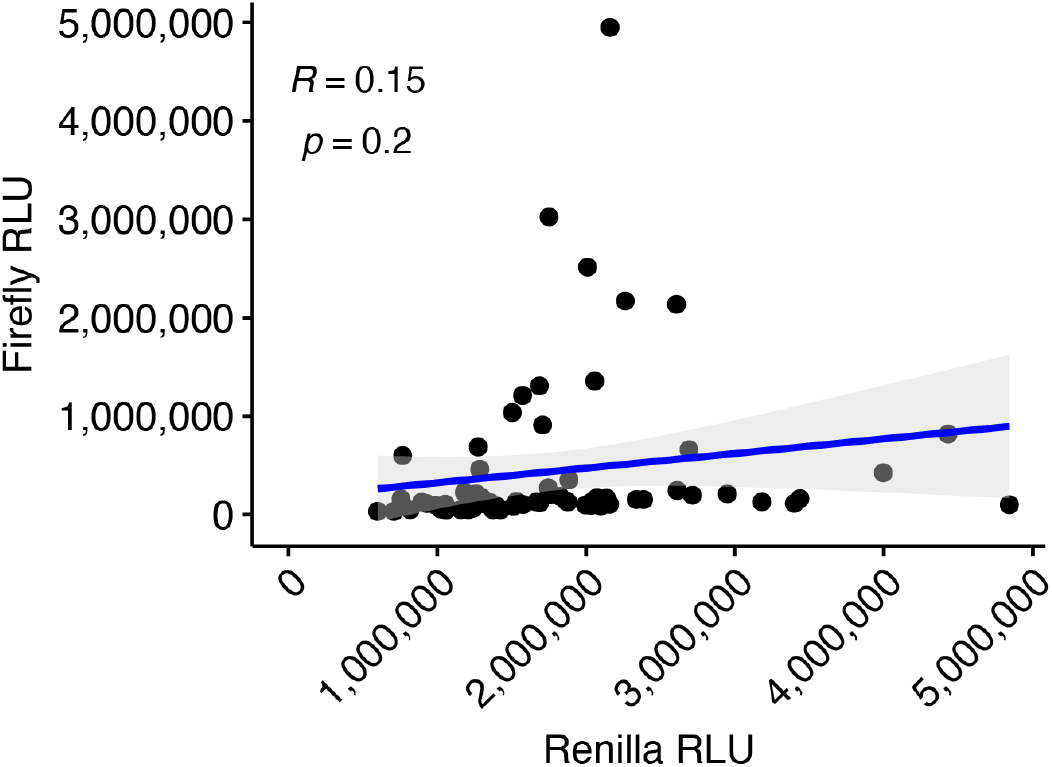
*35S* Batch One *S. viridis* Dual Luciferase Assay. The first protoplast isolation for *S. viridis* transformed with the library of scFv-ADs and the single-plasmid dual luciferase vector. Renilla luciferase is driven by a *35S* promoter. Data was fit to a linear regression model, with R and p values shown above.

## Supplementary Note 11: Supplementary Figure 3

**Supplemental Figure 3.**
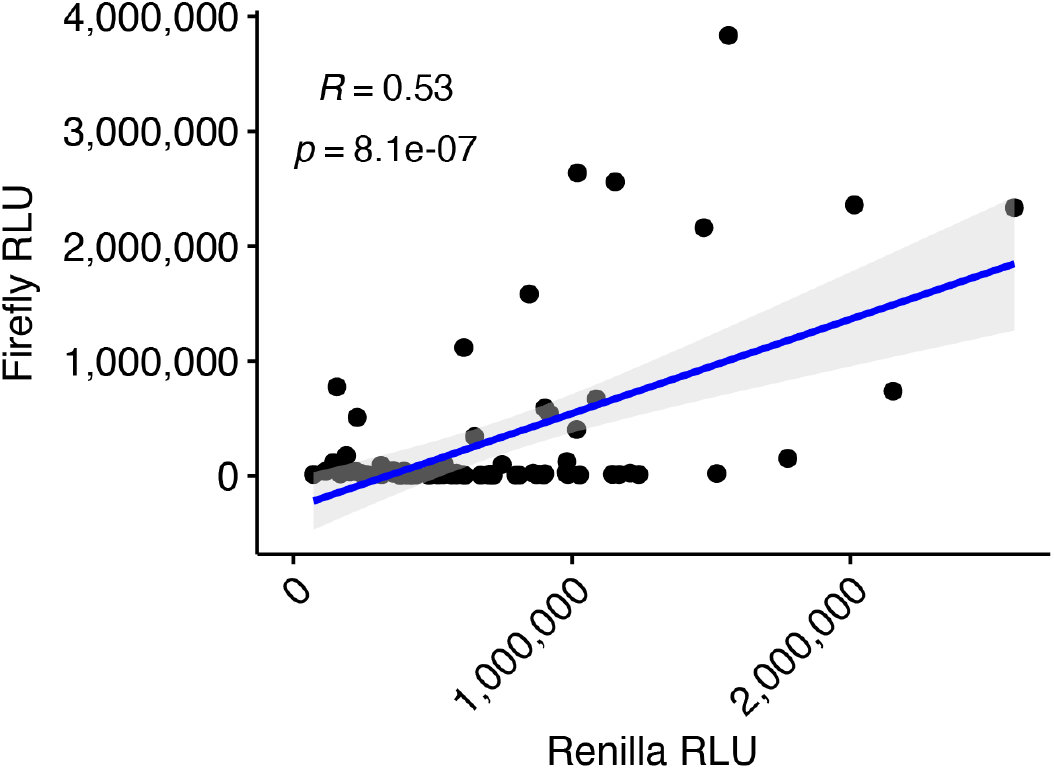
*35S* Batch Two *S. viridis* Dual Luciferase Assay. The second protoplast isolation for *S. viridis* transformed with the library of scFv-ADs and the single-plasmid dual luciferase vector. Renilla luciferase is driven by a *35S* promoter. Data was fit to a linear regression model, with R and p values shown above.

## Supplementary Note 12: Supplementary Figure 4

**Supplemental Figure 4.**
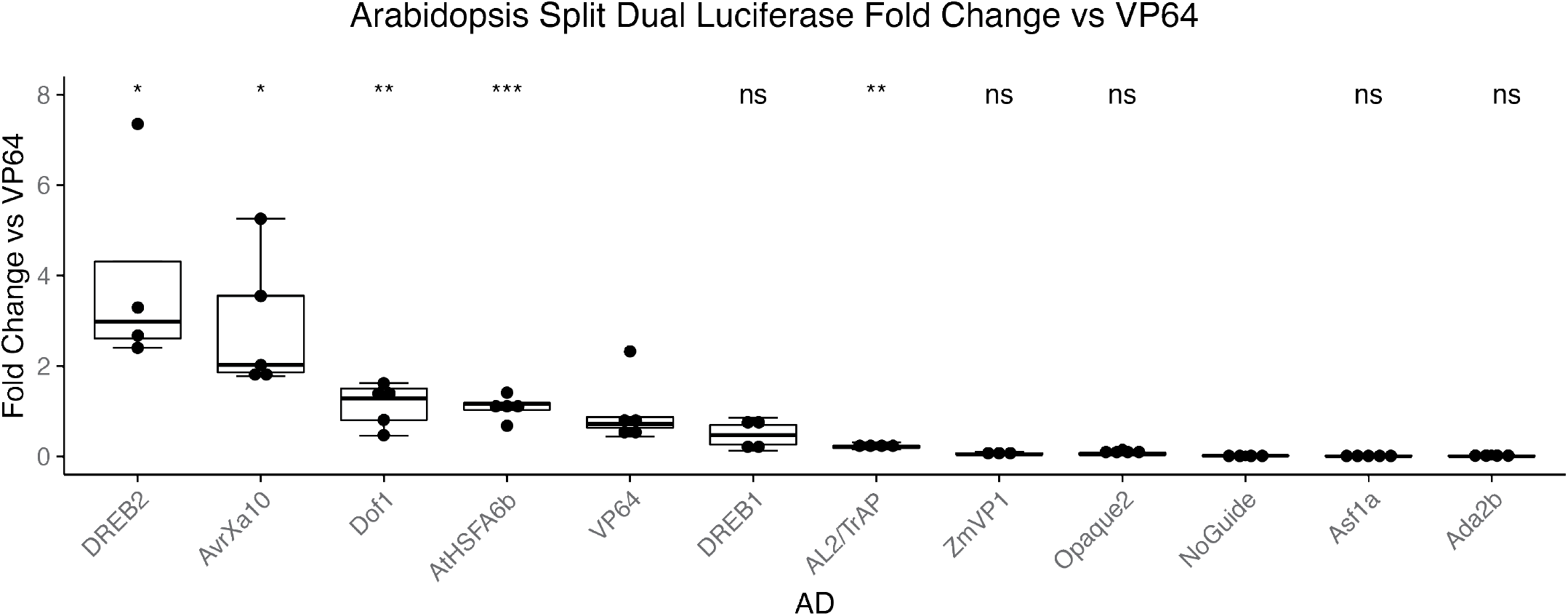
*A. thaliana* Split Dual Luciferase Assay. A smaller library of scFv-ADs was transformed into *A. thaliana* protoplasts along with the split dual luciferase vectors. The firefly luciferase vector contained the *6xLacO-mini35S* promoter, while the Renilla luciferase vector was driven by the constitutive *35S* promoter. Firefly luciferase RLUs were divided by Renilla luciferase RLUs to normalize for transformation. The scFv-AD normalized RLUs were then divided by the scFv-VP64 normalized RLU to generate the plot show above.

## Supplementary Note 13: Supplementary Figure 5

**Supplemental Figure 5.**
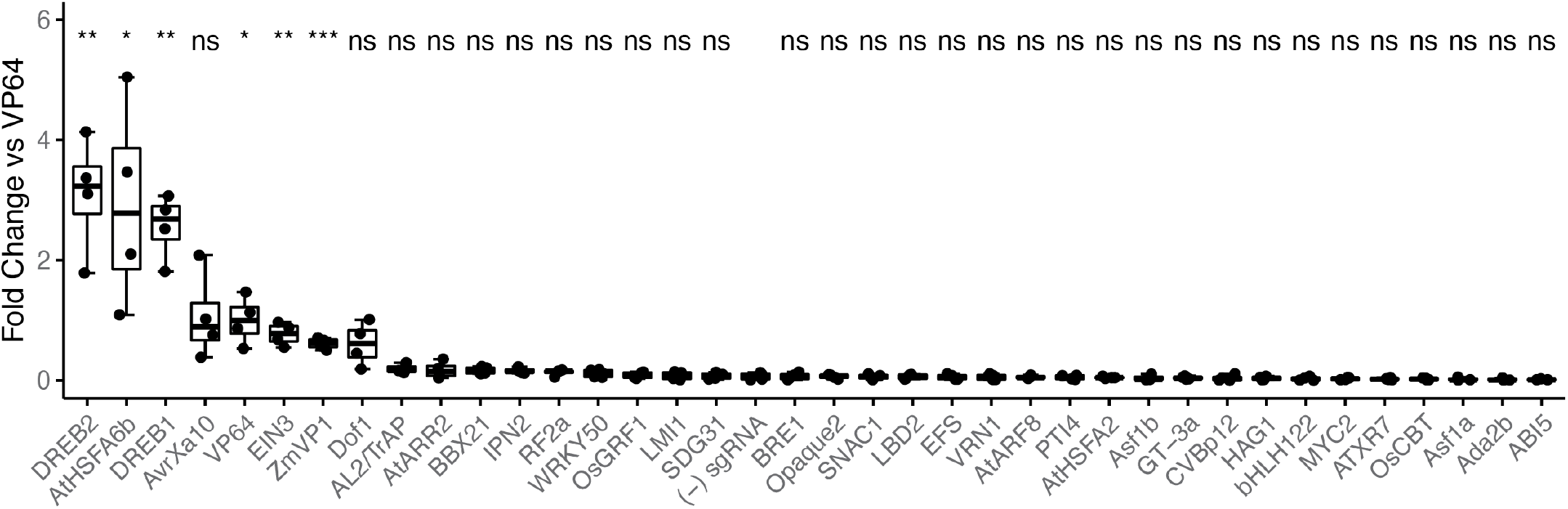
*S. viridis* Single-Plasmid Dual Luciferase Assay. The full library of scFv-ADs was transformed into *S. viridis* protoplasts along with the single-plasmid dual luciferase vectors. The single luciferase reporter contained the firefly luciferase driven by the *6xLacO-mini35S* promoter, while the Renilla luciferase was driven by the constitutive *35S* promoter. Firefly luciferase RLUs were compared to the positive control VP64. A parametric Welch’s t-test was performed comparing the negative control (-) sgRNA with each respective AD. p <= 0.05, ** corresponds to p <= 0.01, *** corresponds to p <=0.001, and **** corresponds to p<= 0.0001.

## Supplementary Note 14: Supplementary Figure 6

**Supplemental Figure 6.**
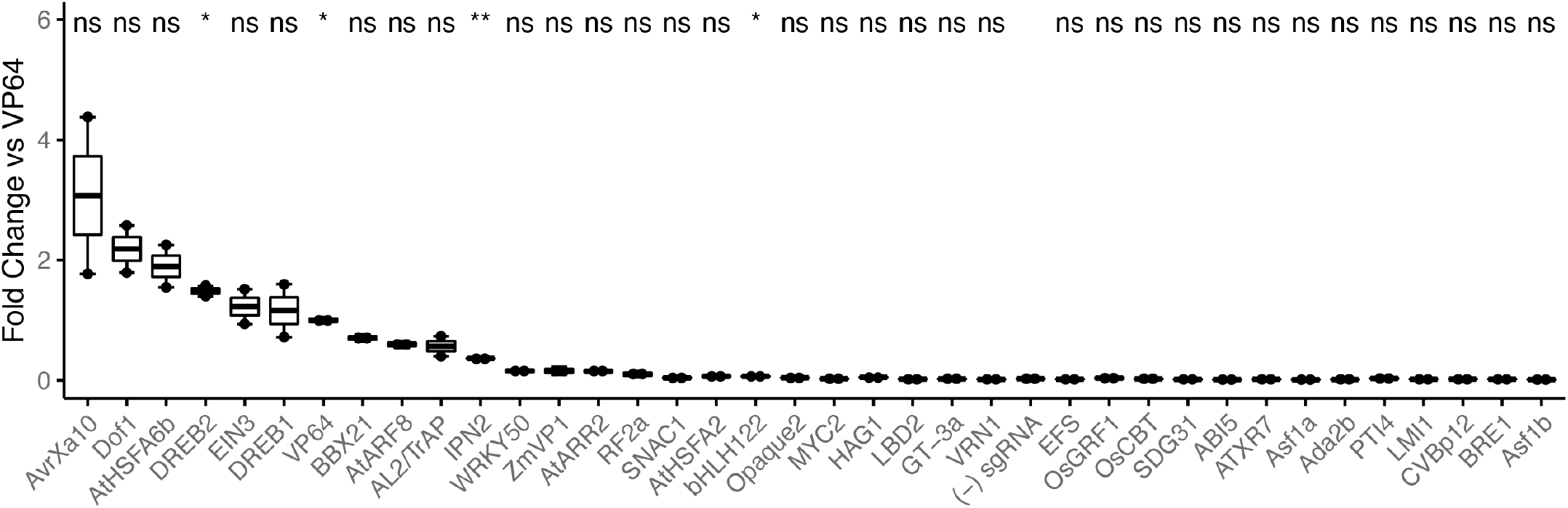
*A. thaliana* Single-Plasmid Dual Luciferase Assay. The full library of scFv-ADs was transformed into *A. thaliana* protoplasts along with the single-plasmid dual luciferase vectors. The single luciferase reporter contained the firefly luciferase driven by the *6xLacO-mini35S* promoter, while the Renilla luciferase was driven by the constitutive *35S* promoter. Firefly luciferase RLUs were compared to the positive control VP64. A parametric Welch’s t-test was performed comparing the negative control (-) sgRNA with each respective AD. p <= 0.05, ** corresponds to p <= 0.01, *** corresponds to p <=0.001, and **** corresponds to p<= 0.0001.

## Supplementary Note 15: Supplementary Figure 7

**Supplemental Figure 7.**
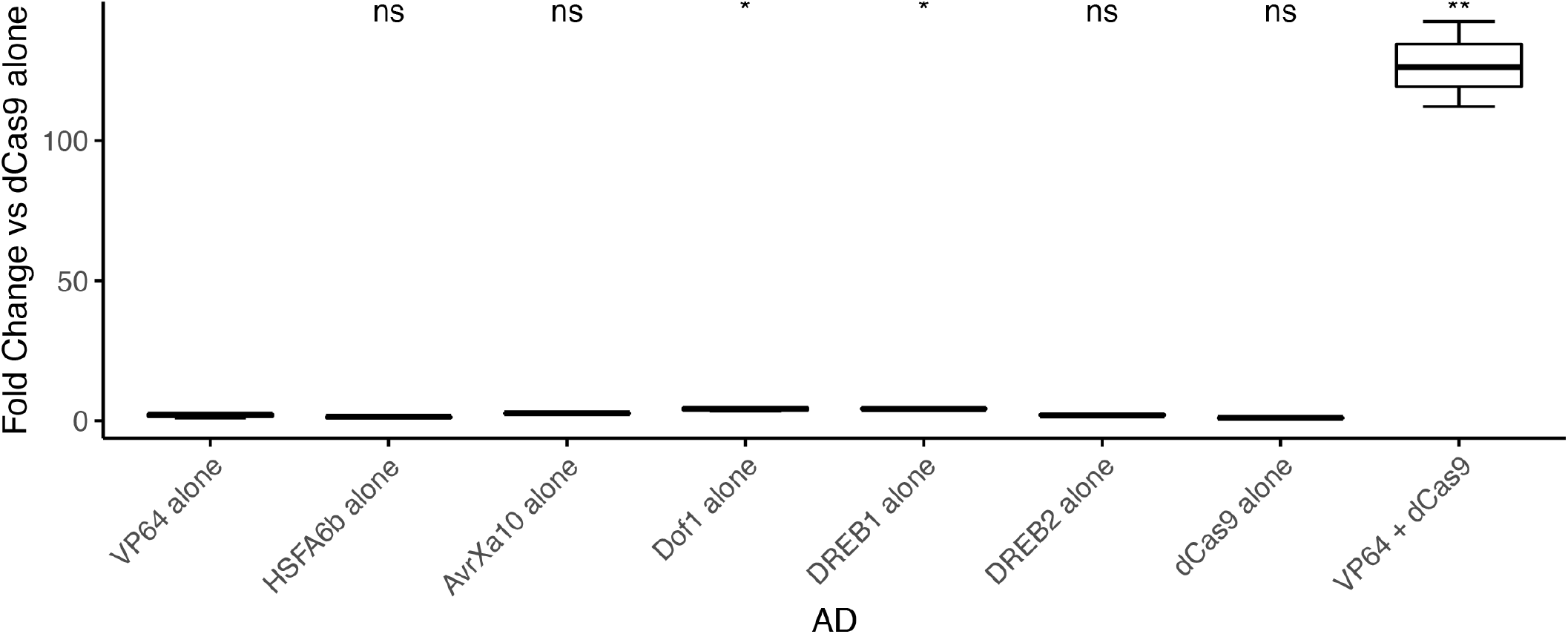
Non-specific Activation Domain strength in *S. viridis* protoplasts. The non-specific activity of each strong plant-derived AD was measured by transforming *S. viridis* protoplasts with the indicated scFv-AD fusion and dual luciferase reporter plasmid. A parametric Welch’s t-test was performed to compare the negative control ‘VP64 alone’ with each individual scFv-AD treatment. The positive control of ‘VP64+ dCas9’ serves as the positive control, in which the dual luciferase, sgRNA, dCas9, and scFv-VP64 were co-transformed into *S. viridis* protoplasts. A parametric Welch’s t-test was performed comparing the negative control of ‘VP64 alone’ with each respective AD. * corresponds to p <= 0.05, ** corresponds to p <= 0.01, *** corresponds to p <=0.001, and **** corresponds to p<= 0.0001.

## Supplementary Note 16: Supplementary Figure 8

**Supplemental Figure 8.**
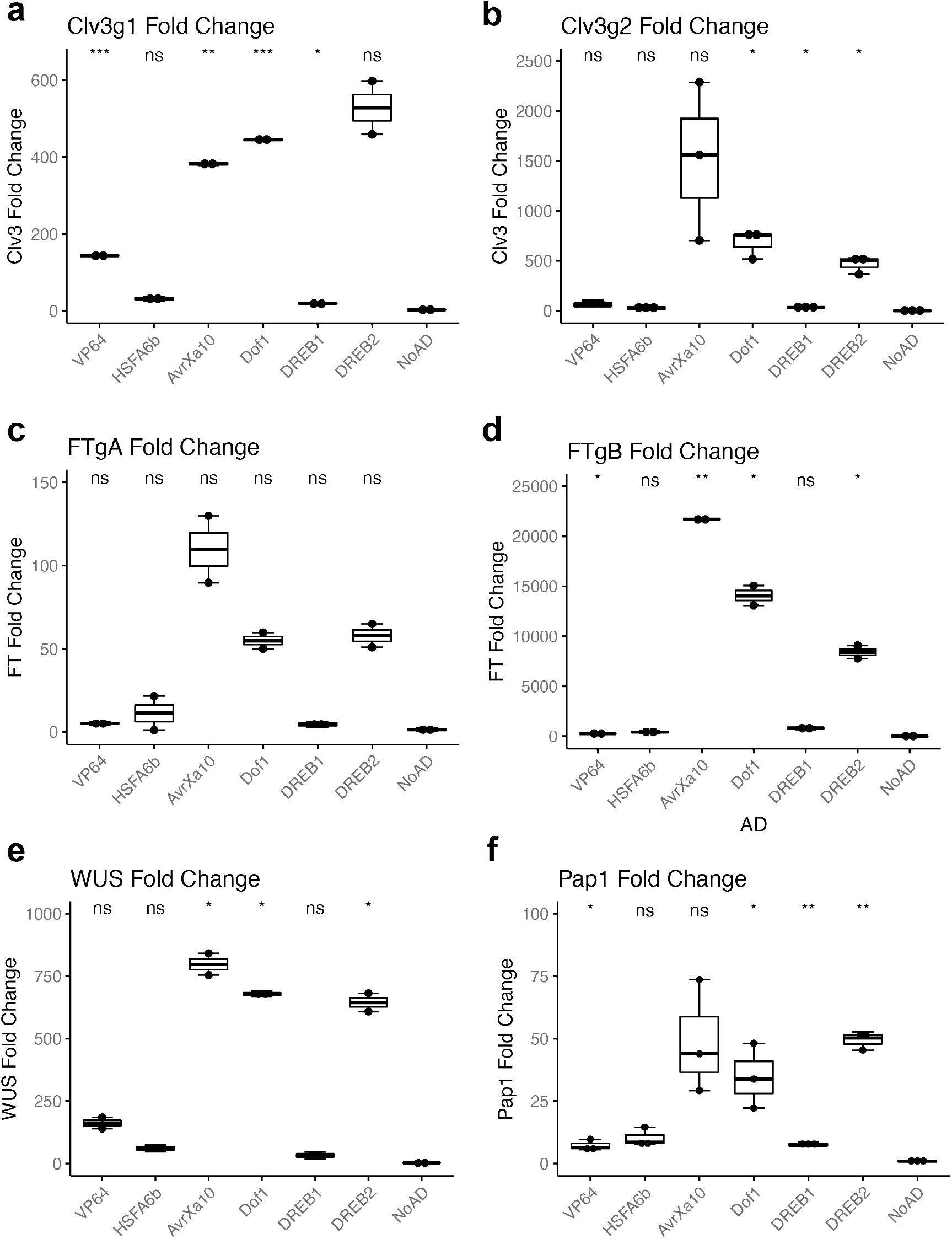
*A. thaliana* protoplast endogenous gene activation across all sgRNAs tested. Protoplasts were isolated from *A. thaliana* and transformed with Ubi10-dCas9-24xGCN, *35S*-scFv-AD, and AtU6-sgRNA constructs for the indicated target gene. RNA was isolated from cells 24h post-transformation, and RT-qPCR was performed to quantify gene expression. The delta delta Ct method was used to calculate Fold Change vs the NoAD negative control. *PP2A* was used as the housekeeping gene. Parametric Welch’s t-tests to compare each AD with the NoAD control are denoted by asterisks, where * corresponds to p<= 0.05, ** corresponds to p<=0.01, *** corresponds to p<= 0.001, and **** corresponds to p<=0.0001.

## Supplementary Note 17: Supplementary Figure 9

**Supplemental Figure 9.**
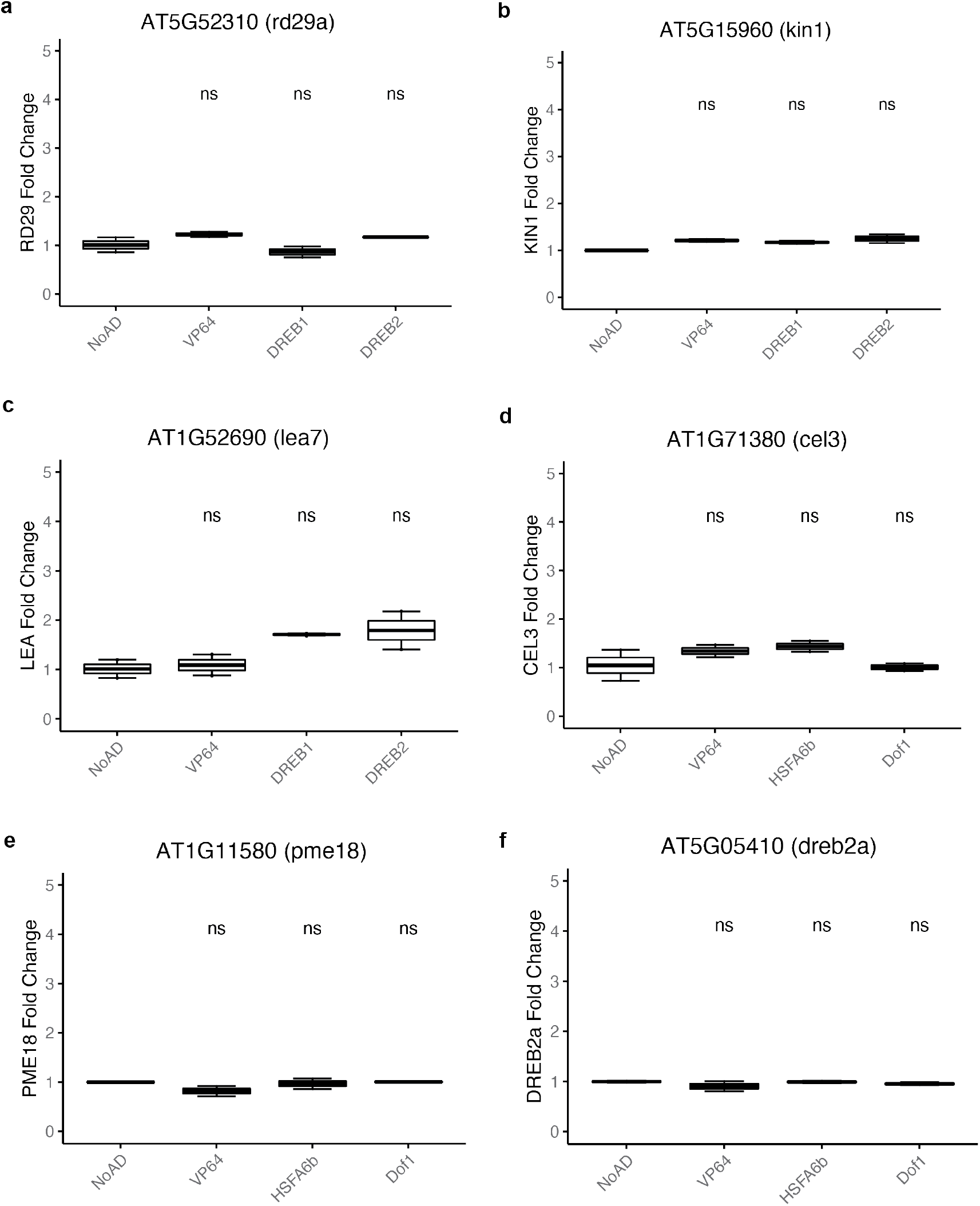
Off-target gene activation in *A. thaliana* protoplasts. The off-target activity of strong plant-derived activation domains was quantified using RT-qPCR for downstream target genes of the endogenous transcription factors. A parametric Welch’s t-test was performed to compare downstream gene expression for a given AD with the negative control “NoAD” in which protoplasts were transformed with dCas9-24xGCN4 and sgRNA but lacking the AD plasmid. Downstream target genes were identified by a literature search. The on-target guide in these experiments was the sgRNA targeting the WUS promoter region.

## Supplementary Note 18: Supplementary Figure 10

**Supplemental Figure 10.**
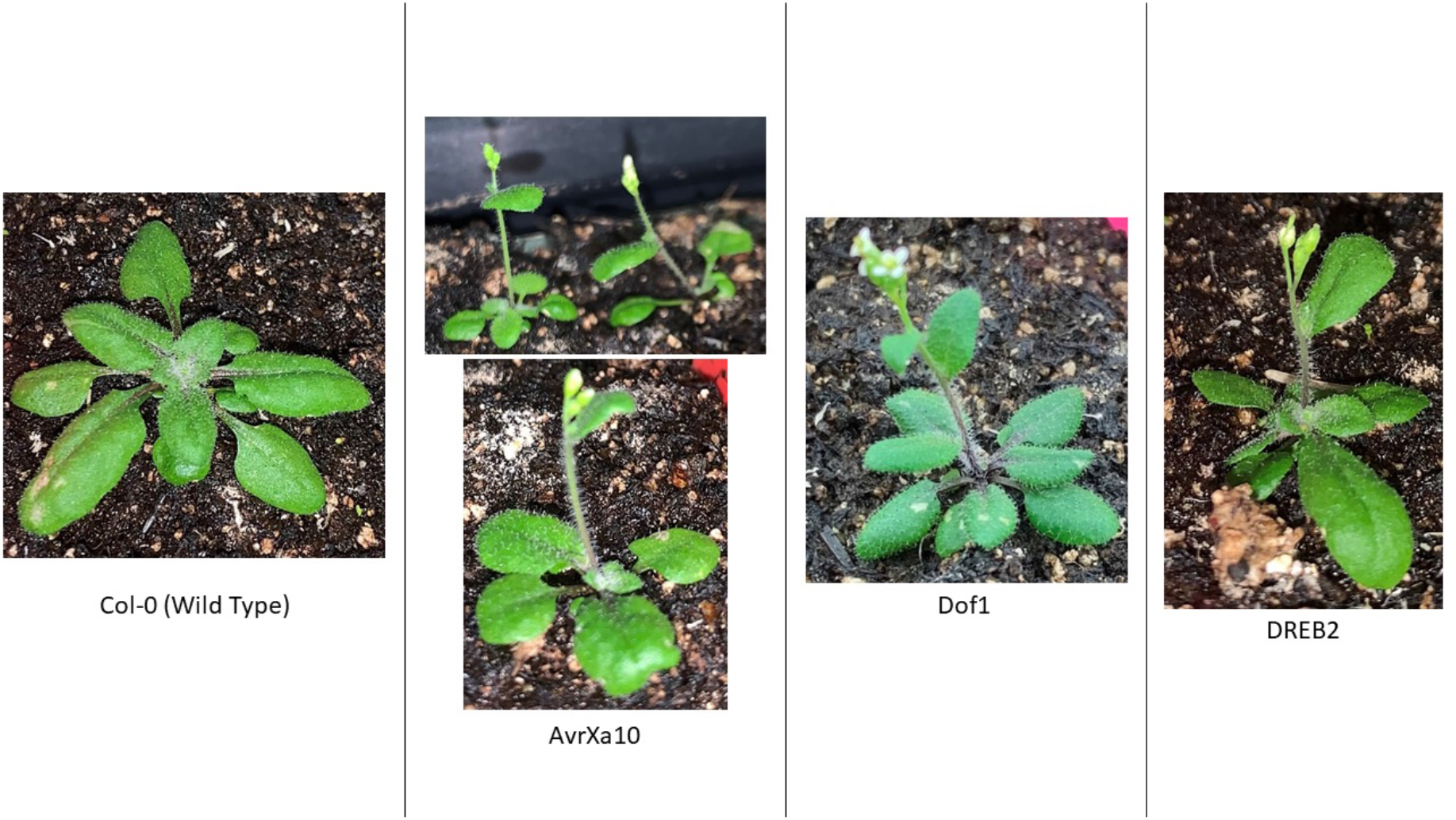
T1 *A. thaliana* Parental Lines. Images showing the T1 parental lines and an age-matched wild type Col-0 *A. thaliana* plant. The binary vectors for these plants are identical, except for the AD fused to NbGP41. Labels below each image denote the AD in each parental line. While a Col-0 line was used as a negative control side-by-side with the FT overexpression lines in this experiment, we observed the same phenotype with an age-matched no-sgRNA control(37).

## Supplementary Note 19: Supplementary Figure 11

**Supplemental Figure 11.**
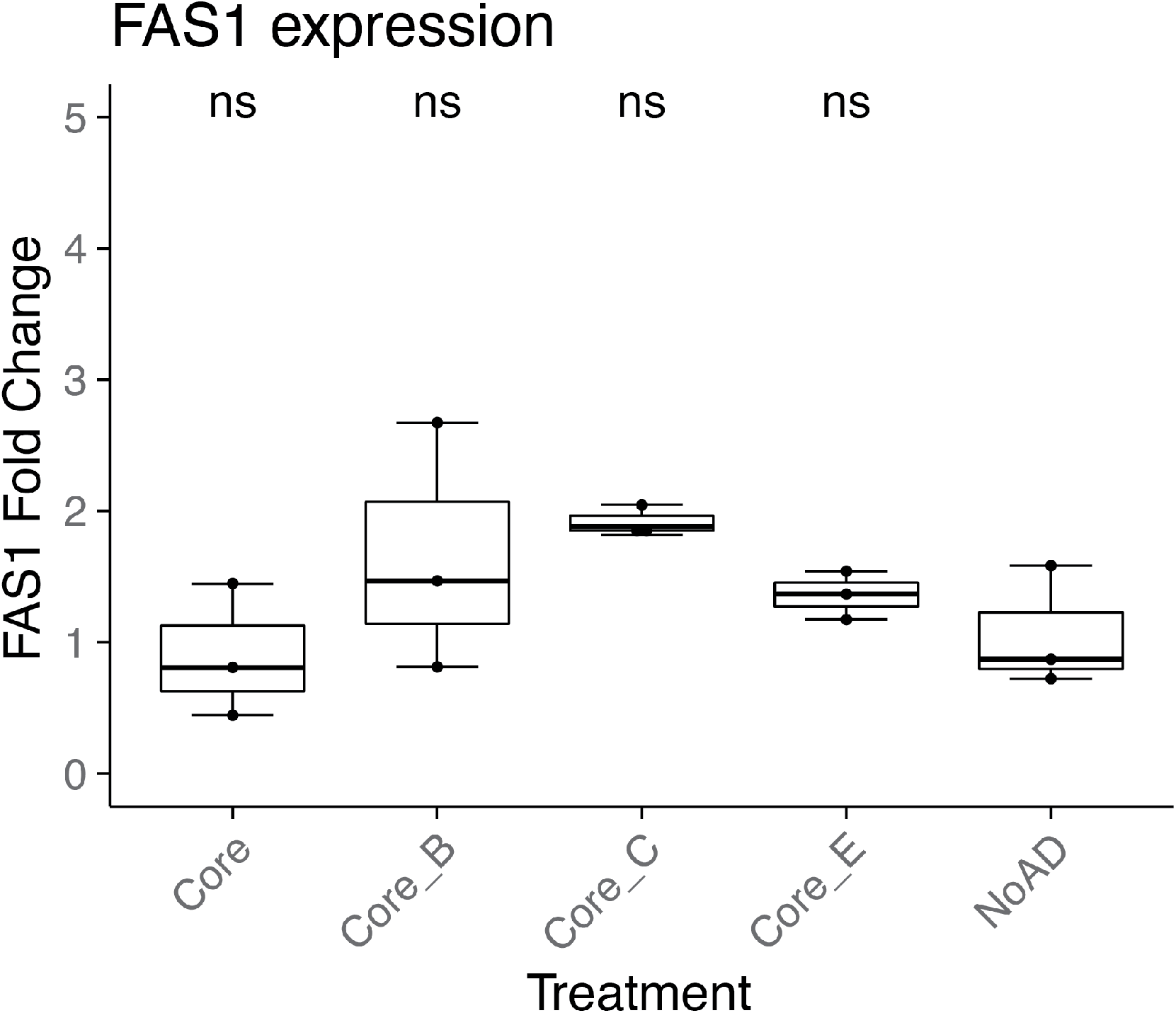
Off-target activation of nearby FAS1 gene in A. thaliana protoplasts. The off-target activity of FAS1 (AT1G65470) was quantified by RT-qPCR in *A. thaliana* protoplasts. The on-target guide in these experiments are denoted below, in which the FT locus was targeted with sgRNAs to activate both the core promoter and nearby enhancers Block B, C, and E as described in other figures. Little FAS1 expression change was measured when comparing across groups using a parametric Welch’s t-test to compare each treatment with the FT Core + No AD negative control.

